# Peptide nanoparticles for systemic mRNA delivery in rodents and non-human primates

**DOI:** 10.1101/2025.11.26.690657

**Authors:** Tõnis Lehto, Helena Sork, Annely Lorents, Julia Rädler, Safa Bazaz, Samantha Roudi, Oskar Gustafsson, Obedoulaye Boukary, Iris R Talgre, Mattias Hällbrink, Oscar P.B. Wiklander, Osama Saher, Jeremy Bost, Kariem Ezzat, CI Edvard Smith, Dhanu Gupta, Taavi Lehto, Samir EL Andaloussi

## Abstract

The therapeutic potential of mRNA is vast, and yet translating this potential into effective treatments requires overcoming significant challenges of achieving safe and efficient delivery. This process is hampered by biological barriers that limit cellular uptake, degrade exposed mRNA, and thus necessitate effective endosomal escape to reach the cytoplasm. To address these challenges, we developed a hPep peptide-based nanoparticle (PNP) system that encapsulates mRNA, forming stable and biocompatible particles, which are rapidly taken up by the cells and enable efficient mRNA delivery across various cell culture models. Following systemic administration in mice, lead hPep3/mRNA PNPs achieve broad mRNA expression across multiple tissues, including the lungs, liver, and spleen, and enable effective mRNA delivery to the central nervous system upon local administration. Furthermore, we established a high-yield (≥70%) microfluidics-based protocol to scale up the production of well-defined, sterile hPep3/mRNA PNP formulations (approximately 70 nm, PDI around 0.170). Most importantly, in a proof-of-concept study in nonhuman primates (NHPs), we demonstrate that hPep PNPs loaded with human erythropoietin (hEPO) mRNA induce dose-dependent expression of hEPO protein in monkey serum, reaching up to 10 ng/ml at 1.0 mg/kg dose, following both single and repeated administration, while remaining systemically well-tolerated. These findings underscore the potential of hPep peptide-based nanoparticles as a versatile platform for mRNA delivery across multiple tissues, highlighting their promise in advancing the development of mRNA therapeutics.

## Introduction

mRNA-based therapeutics have revolutionized medicine, offering versatility for precision therapies [1]. Their adaptability enables applications ranging from personalized cancer vaccines [2,3] and in vivo CAR T-cell generation [4] to treatments for genetic disorders [5]. In oncology, mRNA vaccines can train the immune system to recognize tumor-specific antigens, enabling personalized approaches [6]. The rapid development of COVID-19 vaccines by Pfizer-BioNTech and Moderna further highlighted mRNA’s agility in combating emerging infectious diseases and genome editing applications [7]. Additionally, mRNA enables targeted protein delivery, with ongoing studies in conditions like cystic fibrosis [8] and propionic acidemia [9].

Despite these advances, effective delivery beyond the liver remains a challenge. mRNA is prone to enzymatic degradation and has poor cellular uptake due to its large size, negative charge and hydrophilicity [1]. Encapsulation, commonly with lipid nanoparticles (LNPs), protects mRNA and facilitates liver-targeted delivery [10]. However, achieving efficient extrahepatic delivery to poorly fenestrated organs is limited by the complexity of LNPs and organ-specific targeting hurdles [11,12].

Other non-viral delivery systems are emerging as alternatives, offering flexible designs to reach diverse tissues. These include polymeric nanoparticles [13], cationic polymers [14], and cell-penetrating peptides (CPPs) [15]. CPPs are cationic and/or hydrophobic peptides that can transport diverse nucleic acid cargo across cellular membranes [15,16] — including antisense oligonucleotides, siRNAs, plasmids, and mRNAs — either by direct conjugation or as nanoparticles. Their broad biodistribution enables delivery not only to the liver, but also to the lungs, spleen, muscles, heart and the bone marrow [17–20]. In the case of nanoparticle-based approaches, CPPs, like cationic polymers, leverage their cationic nature to electrostatically encapsulate negatively charged nucleic acids into stable nanoparticles [21]. However, bloodstream instability and reduced membrane interactions – due to competing charged molecules and the inherent hydrophilicity of classical CPPs – still limit delivery efficiency.

To overcome these limitations, structural modifications to CPPs have been pursued to improve stability, cellular uptake, and endosomal escape. Strategies such as peptide cyclization [22], incorporation of endosomal escape-promoting agents [18], and lipid tail conjugation [23] have shown promise, with lipid-modified CPPs, like the PepFect family, demonstrating enhanced stability, nanoparticle hydrophobicity, and interactions with cell membranes [18,24,25].

Building on this, we have developed a new class of hydrophobically modified CPPs (hPeps) by incorporating alkenyl side chains along the peptide backbone [26,27]. This design enables fine-tuning of the hydrophobicity and charge distribution, offering a distinct approach compared to terminal lipid conjugation [28]. Here, we demonstrated that hPeps, systematically varied in alkenyl side chain number and length, form stable mRNA-loaded PNPs. These PNPs rapidly enter cells via endocytic pathways and efficiently escape endo-lysosomal compartments, and release mRNA for robust cytoplasmic translation across diverse cell types.

Systemic administration of PNPs in mice resulted in dose-dependent and well-tolerated delivery of modified firefly luciferase (FLuc) mRNA, driving strong expression across lungs, liver, and spleen. Localized delivery via intracerebroventricular (ICV) injection further demonstrated the platform’s versatility, achieving robust protein expression within the CNS. To support scalable and reproducible manufacturing, we developed a microfluidics-based formulation process that produces monodisperse, sterile PNPs with reduced serum protein adsorption, translating to enhanced in vivo performance. Importantly, proof-of-concept studies in NHPs showed that hPep PNPs loaded with hEPO mRNA mediated a consistent, dose-dependent increase in serum hEPO following both single and repeated administration, with acceptable tolerability. Together, these findings establish the hPep platform as a robust and scalable solution for therapeutic mRNA delivery.

## Results

### hPep peptides form stable nanoparticle formulations with mRNA and mediate cellular uptake via endocytosis

CPPs have been widely studied for nucleic acid delivery [18,29] and are typically cationic and/or amphipathic. The hPeps were designed as α-helices with lysines aligned on one face and leucines on the other, with additional hydrophobic alkenyl-alanines incorporated to increase amphipathicity, thereby strengthening nucleic acid complexation and cellular uptake (Fig. 1a,b and Supplementary Fig. 1).

**Figure 1.**
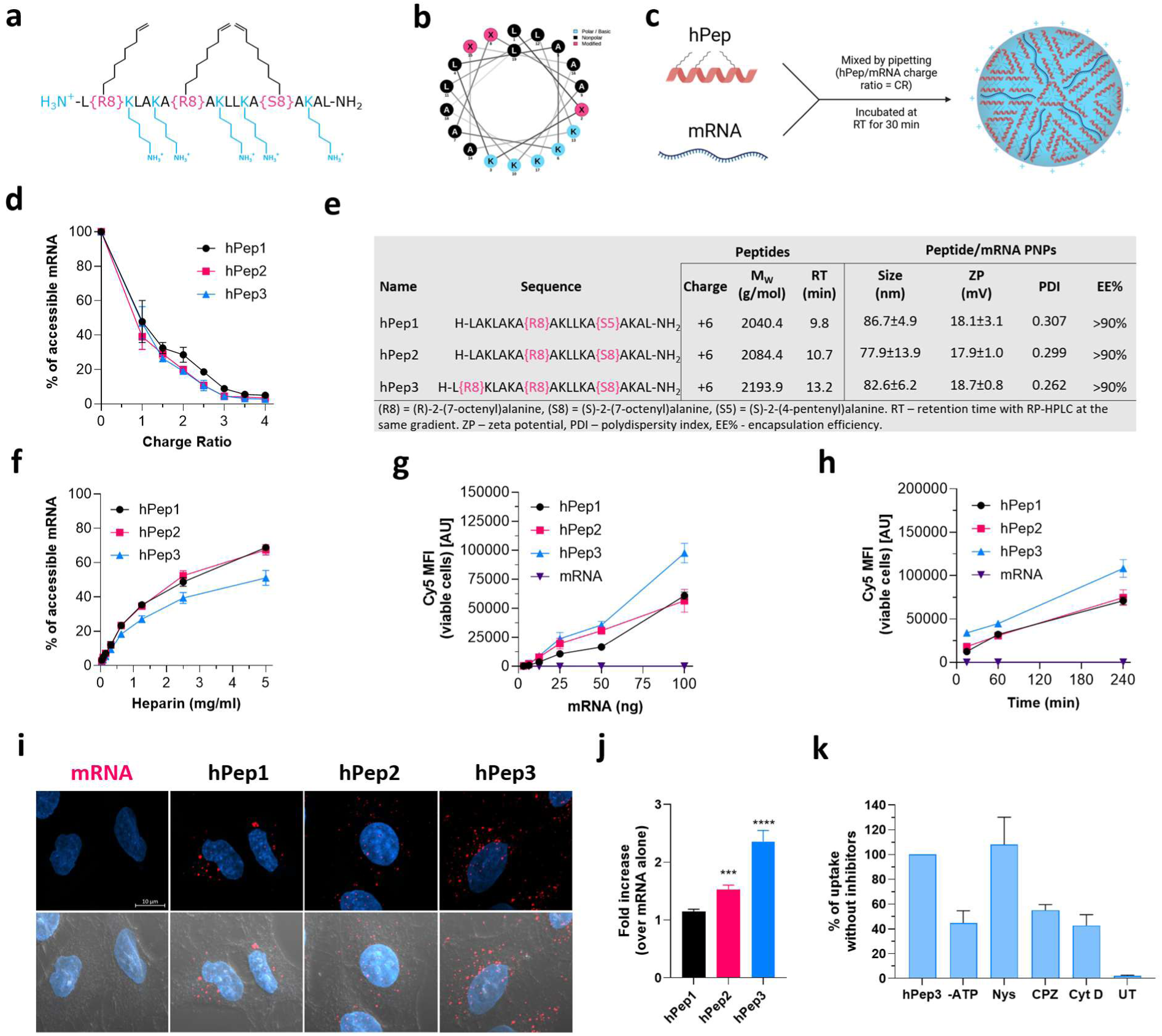
**Design and physicochemical characterization of hPep peptides and peptide/mRNA nanoparticles. a-c**, Schematic representation of hPep3 peptide, its helical wheel projection and non-covalent peptide/mRNA complex formation between positively charged peptide and negatively charged mRNA. **d,** Encapsulation efficiency of hPep/mRNA complexes over charge ratios. **e,** Physicochemical characteristics of hPep peptides and hPep/mRNA PNPs at CR4. **f,** Stability of hPep/mRNA NPs at CR4 to competitive polyanion, heparin, treatment at different concentrations. **g**, Dose-dependent quantitative uptake of hPep/Cy5-mRNA NPs after 4 hours in A549 cells by FACS. **h**, Time-dependent quantitative uptake of hPep/Cy5-mRNA NPs at 50 ng mRNA dose in A549 cells by FACS. Data is presented as mean ± stdev (n=3). **i**, Confocal microscopy of A549 cells treated with Cy5-mRNA NPs at CR4 for 4 h. **j**, Statistical analysis of the uptake of hPep/mRNA NPs (mean ± SEM). Kruskal-Wallis test with Dunn’s multiple comparison. **k,** Uptake inhibition of hPep3/Cy5-mRNA NPs by endocytosis inhibitors (mean ± SEM, n=3).

To evaluate their capacity for mRNA delivery, hPeps were formulated with in vitro transcribed mRNA at increasing peptide-to-mRNA charge ratios (CRs). All hPeps condensed mRNA efficiently, achieving >90% encapsulation at CR4, which was selected as the optimal formulation condition (Fig. 1c,d). At CR4, PNPs were approximately 80 nm in size with a surface charge of +18 mV (Fig. 1e and Supplementary Fig. 2). Heparin competition assay showed that increased peptide hydrophobicity improved resistance to polyanion displacement, with hPep3 exhibiting the highest stability (Fig. 1f).

Cellular uptake was next assessed using Cy5-labeled mRNA. A549 cells showed dose-and time-dependent uptake, with hPep3 PNPs performing the best (Fig. 1g,h and Supplementary Fig. 3). Confocal imaging demonstrated punctate intracellular distribution, consistent with endocytic internalization (Fig. 1i,j). Inhibitor studies confirmed that uptake was ATP-dependent, with strong sensitivity to chlorpromazine and cytochalasin D, indicating clathrin-mediated endocytosis and macropinocytosis involvement, while caveolae inhibition had no effect (Fig. 1k).

Intracellular trafficking analysis showed progressive lysosomal colocalization, with about 40% of PNPs detected in lysosomes by 4 h and roughly 4% of lysosomes containing PNPs (Fig. 2). Galectin-8 reporter assays further demonstrated delayed endosomal escape: few events were visible at 4 h, but punctate recruitment increased markedly by 24 h, indicating gradual endosomal disruption (Supplementary Fig. 4).

**Figure 2.**
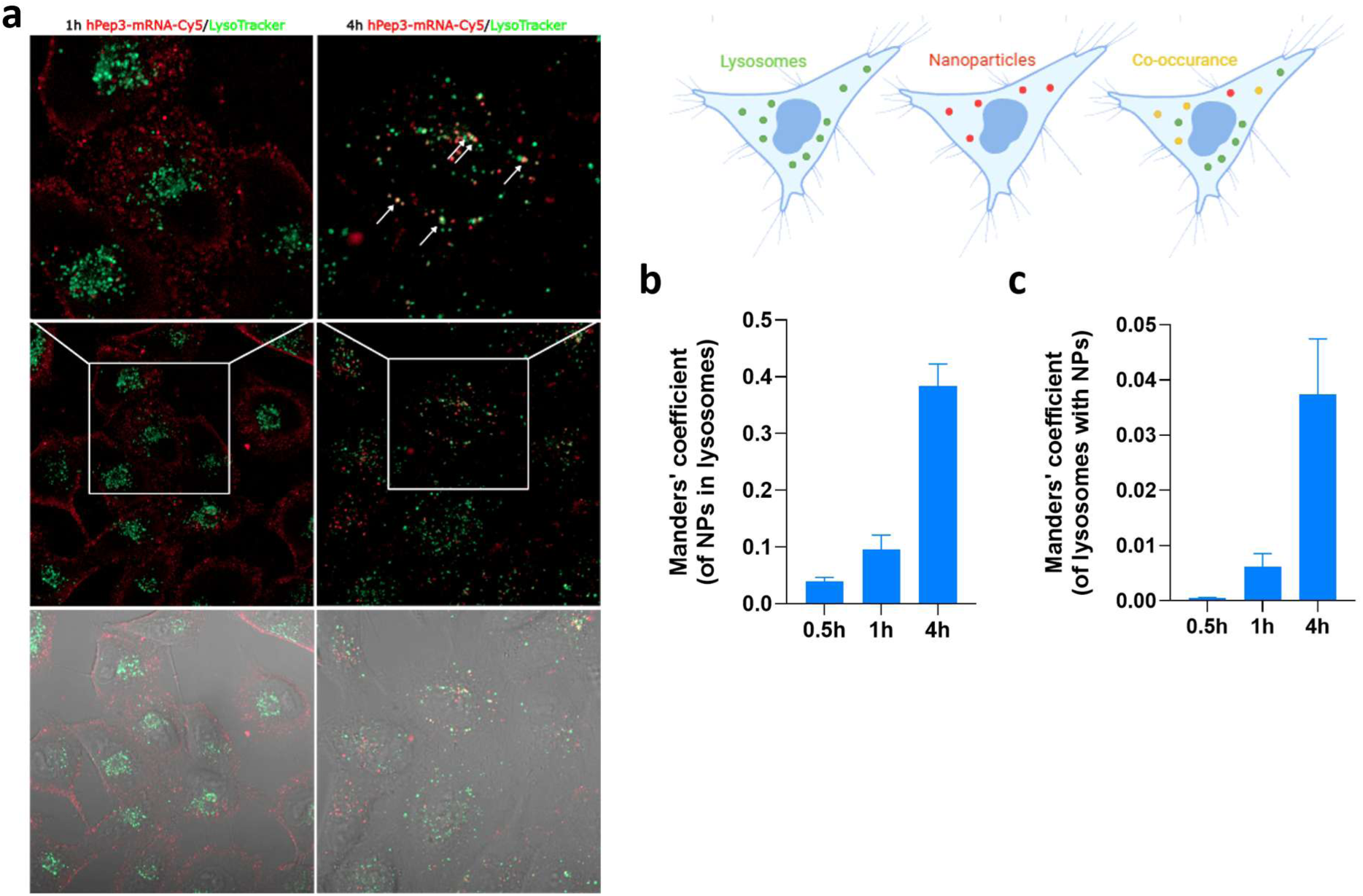
Intracellular trafficking of hPep3/mRNA NPs. a, Intracellular localization of hPep3/Cy5-mRNA NPs and lysosomes by live confocal microscopy was studied in A549 cells at CR4. **b**, Proportion of hPep3/Cy5-mRNA NPs that are trafficked to lysosomes. **c,** Fraction of lysosomes that contain hPep3/Cy5-mRNA NPs. Data is presented as mean ± SEM, n=3.

Together, these data demonstrate that hPeps efficiently condense mRNA into stable nanoparticles with high encapsulation efficiency and stability. The resulting PNPs are readily internalized by cells through multiple energy-dependent endocytic routes, traffic in large extent to lysosomes, and achieve partial endosomal escape to enable mRNA delivery.

### hPep PNPs enable potent, safe, and broad mRNA delivery across diverse cell types

After confirming that hPep peptides can form stable PNPs which are efficiently internalized, we next assessed their bioactivity and safety in cell culture models. PNPs formulated with firefly luciferase (FLuc) mRNA induced robust, dose-dependent protein expression in A549 cells, with increases of up to 4 logs over baseline (Fig. 3a). Interestingly, less hydrophobic derivatives, particularly hPep1, mediated higher luciferase expression than the more hydrophobic hPep3, despite the latter demonstrating the highest uptake (Fig. 1d). Time-course experiments revealed that luciferase activity could be detected at as early as 4 h, peaked at 48 h (Fig. 3b). WST-1 analysis confirmed that the transfections were well tolerated, with minimal impact on cell viability (Supplementary Fig. 5).

**Figure 3.**
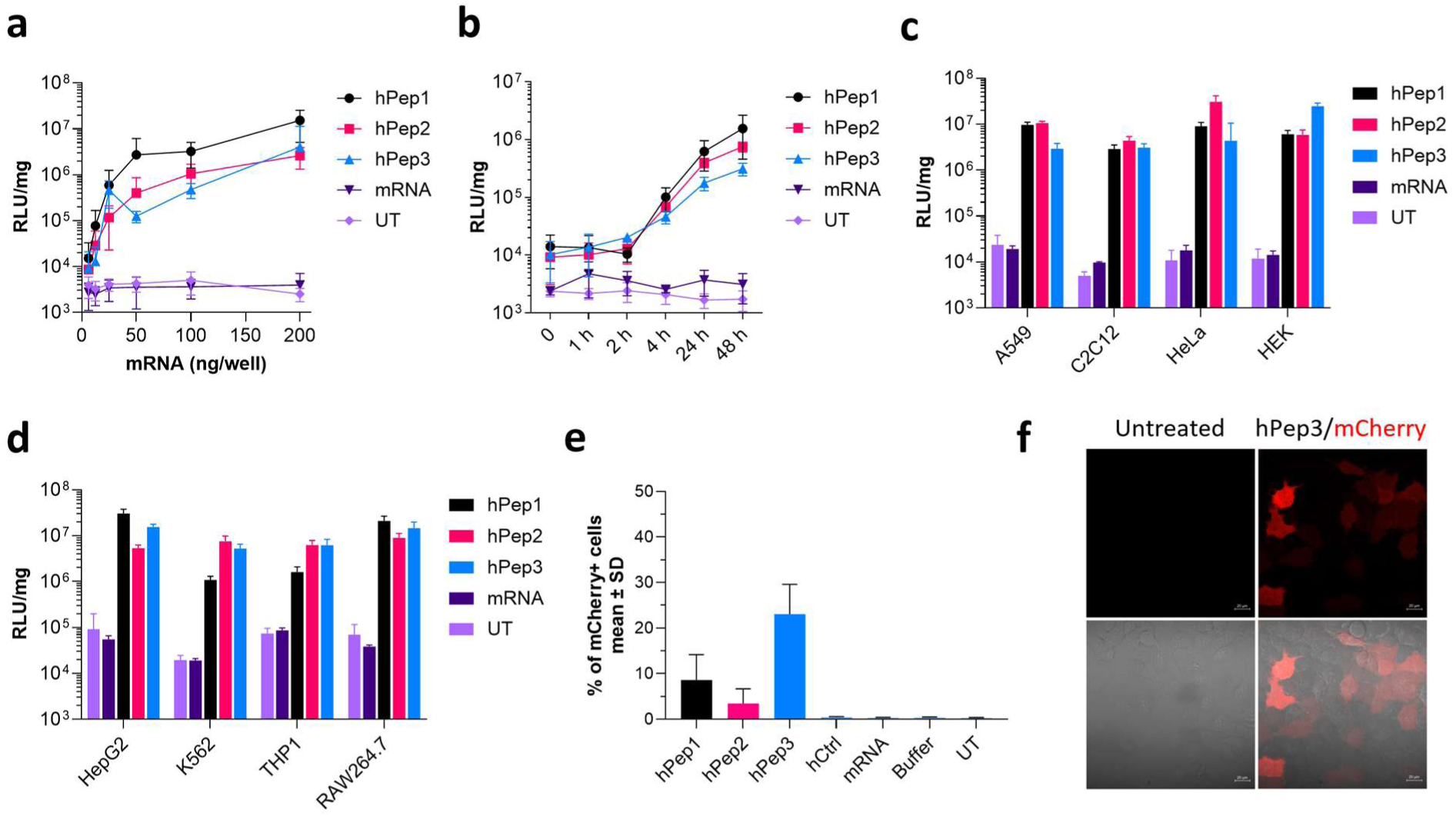
Transfection efficacy in cell cultures. **a**, Dose response of hPep/FLuc-mRNA NPs at CR4 in A549 cells after 24 h. **b**, Transfection efficacy of hPep/FLuc-mRNA NPs (CR4, 100 ng mRNA) in A549 cells under serum-containing conditions at indicated time points. **c-d,** Ability of hPeps to transfect different cell types was evaluated at CR4 at 100 ng dose after 24 h. mCherry expression distribution of hPep3/mCherry-mRNA NPs (CR4) across cell population was evaluated by **e,** FACS, and **f,** confocal microscopy after 24 h. Data is presented as mean ± stdev, n=3.

We next assessed whether the delivery potential extended to other cell types. Bioactivity assays performed across a diverse panel, including myoblasts (C2C12), cervical cancer cells (HeLa), embryonic kidney cells (HEK-293T), hepatocytic cancer cells (HepG2), erythroleukemia cells (K562), monocytic leukemia cell (THP1), and RAW264.7 macrophages, hPeps demonstrated strong mRNA expression efficiency without a clear relationship between peptide hydrophobicity and activity (Fig. 3c,d). Screening across charge ratios revealed that efficacy generally increased with peptide excess, peaking at CR4, though some cell types showed maximal activity at CR2 (Supplementary Fig. 6). This suggests that optimal PNP bioactivity requires both stable particle formation and enhanced membrane interaction, consistent with reduced expression upon endocytosis inhibition (Supplementary Fig. 7).

To test versatility for delivering different mRNA cargos, we formulated hPep PNPs with mCherry mRNA. Flow cytometry analysis confirmed robust protein expression in A549 cells, particularly with hPep3, and confocal microscopy verified widespread distribution (Fig. 3e,f). These results indicate that hPeps can deliver different mRNA cargos across a broad range of cell types without compromising viability.

Finally, we examined the structural basis of hPep activity. Given prior evidence that alkenyl-alanines, especially octenyl-alanines, are key to function [26,27], we generated a library of substituted hPep3 analogs (Supplementary Table 1). To understand the dependency on the alkenyl modifications and to see if the chemical space around the hPep peptides could be further optimized, we designed a small library of new hPep analogs. Replacement of the alkenyl-alanines with hydrophobic/aromatic residues (tyrosine, phenylalanine, tryptophan) or alanine abolished nanoparticle formation (Supplementary Fig. 8a,b). N-terminal lauric acid conjugation restored encapsulation, but activity was reduced relative to hPep3 (Supplementary Fig. 8c). Similarly, substitution with hydrocarbon-modified glycines or fatty acid–modified lysines produced particles but failed to support efficient expression. Increasing hydrophobicity via octanoic acid conjugation also reduced activity and impaired encapsulation. Collectively, these results demonstrate that alkenyl-alanines are indispensable for hPep activity and cannot be easily replaced.

In summary, hPep peptides mediate efficient, safe, and versatile mRNA delivery across multiple cell types, with bioactivity critically dependent on their unique alkenyl-alanine modifications.

### hPep PNPs achieve systemic mRNA delivery with multi-organ expression

To assess the delivery potential of hPep PNPs in vivo, formulations carrying FLuc mRNA were administered via tail vein injection in NMRI mice (1.25 mg/kg). Twenty-four hours post-injection, tissues were harvested and analyzed ex vivo with IVIS bioluminescence imaging. Strong luciferase expression was observed across multiple organs, with particularly high activity in lungs, liver, spleen, and heart (Fig. 4a). Among the peptides, hPep3 displayed the most robust delivery profile, substantially outperforming the less hydrophobic hPep1 and hPep2 derivatives. This contrasted with in vitro results where hPep1 and hPep2 displayed equal or higher activity (Fig. 1c), suggesting that in vivo stability—enhanced by hydrophobicity—is a key determinant of functional delivery.

**Figure 4.**
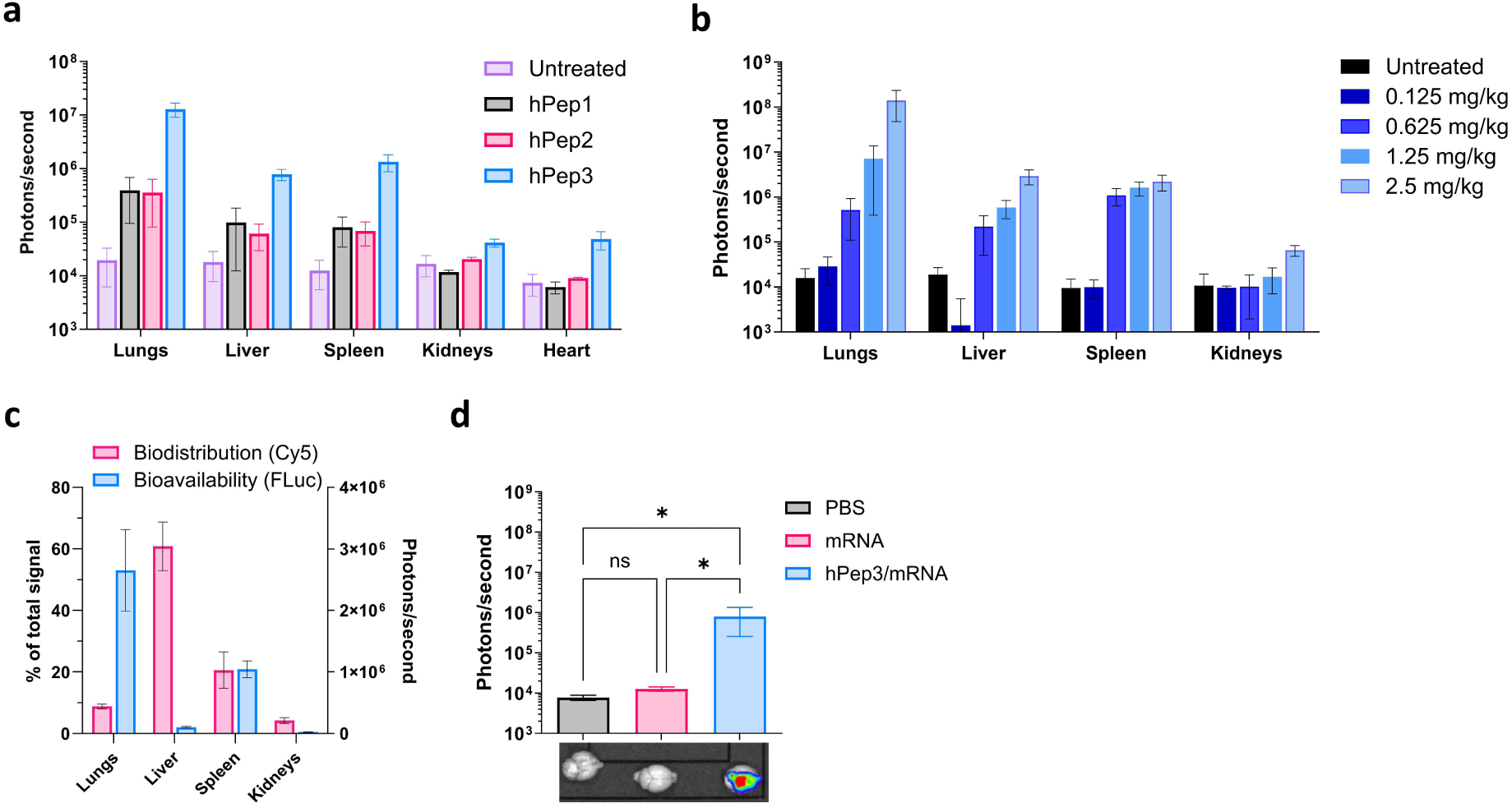
In vivo comparison of hPep/mRNA NPs. a, Comparison of hPeps at i.v. administration of 1.25 mg/kg FLuc mRNA. Luciferase expression is measured with IVIS after 24 h (n=3 animals per group). **b,** hPep3/FLuc mRNA NPs formulated at CR4 were administered i.v. at noted doses. Luciferase expression was measured after 24 h from tissues ex vivo under IVIS (n=3 animals per group). **c,** In vivo bioavailability (Fluc mRNA expression) and biodistribution (Cy5-mRNA accumulation) were compared side-by-side at 2 mg/kg dose at CR4 and quantified ex vivo under IVIS after 24 h. **d,** hPep3/mRNA NPs mediate efficient local mRNA delivery in the CNS. 1 µg of naked FLuc mRNA or 1 µg of hPep3/FLuc-mRNA NPs at CR4 were injected ICV into the brain and measured ex vivo with IVIS after 24 h (n=3 animals per group). Data is presented as mean ± stdev. Statistics: ordinary one-way ANOVA with Holm-Šídák’s multiple comparisons test or two-way ANOVA with Tukey’s multiple comparisons test.

To gain more insight into the structural requirements for in vivo efficacy, we tested NP-forming hPep analogs in which alkenyl-alanines were replaced. From these derivatives, laurylated analogs (C12-hPep3-YFF, C12-hPep3-YWW) could be safely administered but generated only weak luciferase signals. Peptides substituted with hydrocarbon-modified glycines (hPep3-OctG, hPep3-2Ado) displayed moderate activity, mainly in lungs and spleen, exceeding hPep1 and hPep2 but remaining less effective than hPep3. All other hPep variants lacked detectable activity (Supplementary Fig. 8d,e). Together with in vitro data, these results underscore the essential role of alkenyl-alanines for efficient delivery and identified hPep3 as the lead candidate for continued in vivo evaluation.

Dose-escalation studies revealed dose-dependent increase in protein expression across tissues, with dose up to 2.5 mg/kg being well-tolerated (Fig. 4b). At lower doses (e.g. 0.625 mg/kg), expression in lungs dropped sharply, while expression in spleen was preserved, resulting in a more balanced distribution between lungs, liver and spleen. This finding highlights that dosing can modulate organ tropism, potentially enabling tailored delivery to different tissues.

Biodistribution analysis with Cy5-labeled FLuc mRNA revealed notable differences between nanoparticle localization and protein output (Fig. 4c). Although, in NMRI mice, only 8% of hPep3 PNPs localized to lungs, this accounted for 70% of total luciferase expression, pointing to highly efficient translation or stability of mRNA in pulmonary tissue. Conversely, the liver accumulated around 60% of particles but yielded just about 2% of protein expression, suggesting inefficient translation or rapid mRNA degradation. Spleen demonstrated a more proportional relationship between localization and activity (close to 20% in each). Notably, in Balb/c mice, luciferase expression was more evenly distributed, 47% in lungs, 43% in spleen, and 8% in liver (Supplementary Fig. 8d), indicating that strain-specific factors may also influence delivery outcomes.

When compared with MC3-based LNPs, LNPs achieved higher absolute expression levels but were strongly liver-tropic, limiting their application in extrahepatic delivery (Supplementary Fig. 9). In contrast, hPep3 PNPs supported broader biodistribution, with strong lung and spleen activity at higher doses (2.5 mg/kg), suggesting complementary advantages over existing LNP systems.

We next profiled cell-specific expression using hPep3 PNPs formulated with EGFP mRNA. Flow cytometry analysis of the lung and spleen revealed low but distinct transgene expression in selected immune and non-hematopoietic subsets (Supplementary Fig. 10a,b). In the spleen, the GFP-positive cells were primarily macrophages and B cells, while in the lungs most (almost 75%) GFP-positive cells were non-hematopoietic, with close to 20% being macrophages. (Supplementary Fig. 10c,d). These data demonstrate that hPep3 PNPs can target both immune and stromal compartments, highlighting their potential for immunomodulatory therapies.

Finally, to overcome the unmet medical need for central nervous system delivery, hPep3 PNPs were injected intracerebroventricularly. As shown in Fig. 4d, luciferase expression was markedly higher and more broadly distributed across the brain compared to naked mRNA. IVIS imaging confirmed a more extensive and intense luminescence signal, establishing that hPep3 effectively enhances mRNA delivery and functional expression in the CNS.

Collectively, these studies establish hPep3 as a versatile in vivo delivery platform with robust activity, especially in critical organs, the ability to modulate tissue tropism through dosing, and applicability to both systemic and CNS delivery. These findings position hPep3 PNPs as a promising candidate for proof-of-concept pre-clinical studies in higher animals, including non-human primates.

### Microfluidic formulation enables scalable production of uniform hPep PNPs

Rodent studies with hPep3/mRNA PNPs demonstrated robust activity following both systemic and local administration. However, to evaluate the translational capacity, proof-of-concept studies in the NHPs are essential. These require substantially larger formulation quantities than can be achieved with conventional small-scale mixing methods. To address this, we established a scalable microfluidic mixing process for the hPep/mRNA PNPs. Microfluidic mixing provides tight control over particle assembly, yielding homogenous nanoparticles with improved reproducibility and reduced batch-to-batch variability, and, has already been validated for LNP-based clinical formulations [30]. To our knowledge, this is the first systematic development of microfluidic methods for peptide-based mRNA nanoparticles.

We first defined critical formulation criteria for NHP studies. Based on preliminary findings, particle size was targeted at <200 nm (preferably <100 nm) with PDI <0.25 to ensure uniformity. As the hPep/mRNA formulations would be administered intravenously, sterility was required, with maximum injection volume limited to 5 ml/kg (IACUC guidelines), restricting the final mRNA concentration to 0.6 mg/ml.

Optimization was performed on NanoAssemblr™ instruments. After testing several process iterations, and optimizing the conditions for each step separately, we arrived to the final hPep3/mRNA PNP formulation process as illustrated in Fig. 5a. Screening revealed that both components could be dissolved in water, but optimal NP formation required a peptide/mRNA weight ratio of 8:1 – a 2-fold increase compared to formulations used in studies above. Furthermore, on NanoAssemblr™ Blaze™ instrument in combination with NxGen 500 cartridge, the optimal parameters were 100 ml/min total flow rate (TFR), 1:1 flow rate ratio (FRR), and input concentrations of 2.4 mg/ml hPep3 and 0.3 mg/ml mRNA, yielding a homogeneous output formulation at 0.15 mg/ml. Subsequent tangential flow filtration (TFF) enabled concentration, buffer exchange, and sterilization, achieving final formulations at 0.6 mg/ml mRNA in 10 mM HEPES, 10% sucrose (pH7.4). Preparations were stable when cryo-stored at –80 °C and remained stable after repeated freeze-thaw cycles (Supplementary Fig. 11). Full parameters of the formulation process are detailed in Supplementary Table 2.

**Figure 5.**
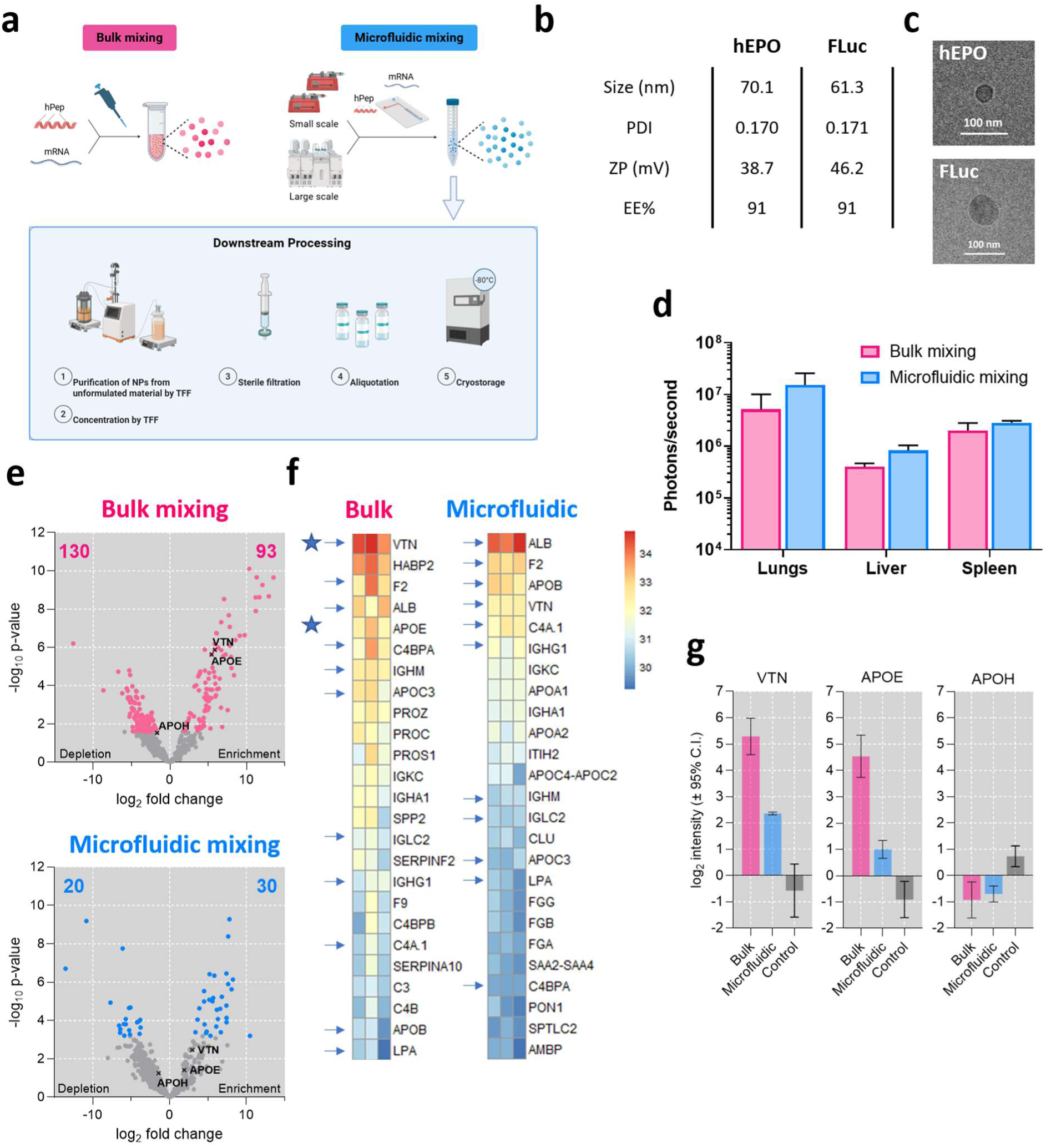
Microfluidics production and scale-up of hPep3/mRNA formulations. a, Overall schematic of the formulation process of hPep/hEPO mRNA nanoparticles. **b-c,** Properties of NP formulations used in the NHP trial by DLS and cryoEM. **d,** In vivo study in mice comparing IV administration of the hPep3/Fluc mRNA bulk and microfluidics NP formulations at 1 mg/kg. The luciferase expression was measured after 24 h (n=3 animals per group). **e-f,** Protein corona analysis, with label-free proteomics comparing the hPep3/mRNA formulations (Ctrl/Fluc) produced by bulk mixing and microfluidics protocols, showing the enrichment of proteins in corona composition. The heatmaps depict normalized LFQ intensity values of the top 25 proteins (ranked by highest average expression in bulk and microfluidically mixed PNPs, respectively). **g,** Enrichment of vitronectin (VTN), Apolipoprotein E (APOE) and Apolipoprotein H (APOH) in protein corona of bulk vs microfluidic mixed NPs. Relative protein level across samples using log2-transformed protein intensities centered across all samples is depicted. Data is presented as mean ± stdev.

Two formulations were produced for scale-up validation: hPep3/hEPO mRNA and hPep3/FLuc mRNA. Both yielded nanoparticles of approximately 60-70 nm, with PDI <0.2, positive zeta potential and >90% encapsulation efficiency (EE) (Fig. 5b). Cryo-EM confirmed the uniform, spherical morphology (Fig. 5c). The overall recovery of mRNA was >75% for both formulations Supplementary Table 3.

To test whether microfluidic formulations retained functionality, we compared them with bulk-mixed PNPs in mice. hPep3/FLuc mRNA PNPs produced using microfluidics induced stronger luciferase expression while preserving a comparable biodistribution across lungs, liver, and spleen (Fig. 5d), confirming enhanced bioactivity without altering tropism.

Because protein corona formation strongly influences NP biodistribution, we next examined interactions with plasma proteins. Compared to bulk-mixed PNPs, microfluidic formulations showed reduced enrichment of blood proteins, as revealed by volcano plot analysis of the interactome (Fig. 5e). Specifically, apolipoprotein E (APOE) and vitronectin (VTN) were more abundant in the corona of bulk-mixed particles (Fig. 5f,g). These proteins have been linked to altered lung and liver tropism [31], [32], suggesting that the more defined microfluidic formulations interact less promiscuously with blood components, potentially improving targeting and consistency. Notably, ApoH—previously implicated in spleen tropism of LNPs [33] —was absent, despite strong splenic activity observed for hPep3 PNPs, pointing to a distinct biodistribution mechanism.

Together, these results demonstrate the feasibility of producing sterile, homogeneous, and bioactive hPep3/mRNA PNPs at scale using microfluidics. Importantly, reduced protein corona formation underscores the advantage of this approach for achieving reproducible in vivo performance and supports advancement toward NHP studies and eventual clinical translation.

### hPep3 PNPs provide durable and well-tolerated mRNA delivery in non-human primates upon single and repeated administration regimens

Following comprehensive optimization of formulation scale-up and purification, we next evaluated whether the efficacy of hPep3/mRNA PNPs could be translated from rodents to NHPs. As a model therapeutic, we selected hEPO mRNA, which enables simultaneous non-invasive monitoring of efficacy and tolerability, while allowing direct comparison with published LNP-based mRNA delivery studies.

To assess efficacy and tolerability, cynomolgus monkeys were administered with 0.3 mg/kg or 1 mg/kg of hEPO mRNA-loaded hPep3 PNPs via a slow (60-90 s) intravenous bolus injection. Serum EPO levels and full toxicology profiles were monitored over time. hPep3 PNPs induced a clear dose-dependent increase in circulating hEPO, with peak concentrations at 6 h, returning to baseline after 72 h (Fig. 6a). At 1 mg/kg, serum hEPO levels reached over 9700 pg/ml at 6 hours—an >100-fold increase from baseline. This translated into robust rise in reticulocyte count (ARET) and percentage (%RET) (Fig. 6b,c), while RBCs and hematocrit (HCT) remained unchanged (data not shown). Overall, single doses up to 1 mg/kg were well tolerated, with no clinical signs of toxicity.

**Figure 6.**
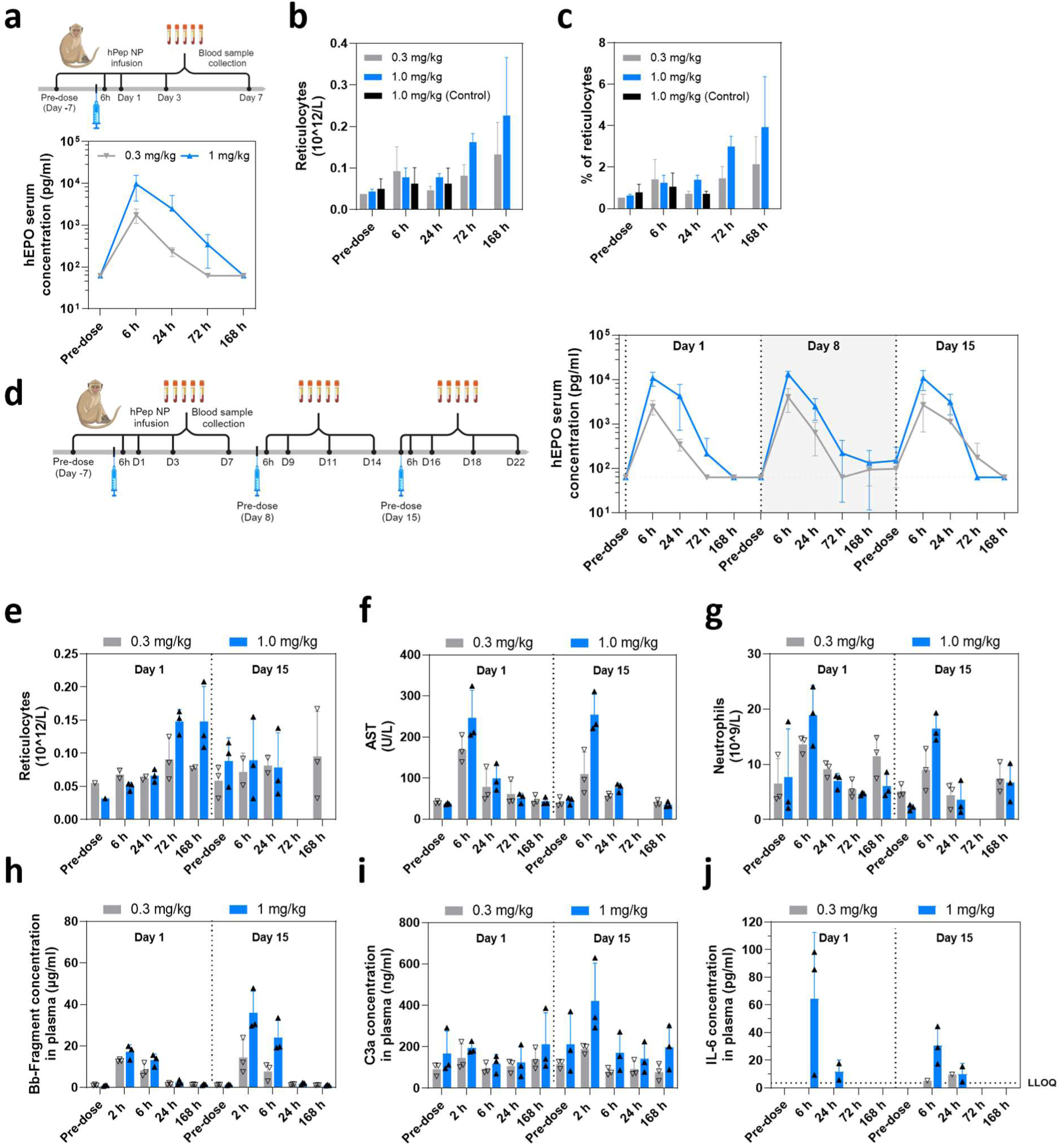
Proof-of-concept trial of hPep3/hEPO mRNA NPs in NHPs. a, Serum hEPO concentrations after single i.v. administration of hEPO mRNA PNP formulations in NHPs (n=3 animals per group) at 0.3 and 1.0 mg/kg dose. **b-c,** Corresponding changes in total reticulocyte count and percentage. **d,** Serum hEPO concentrations after weekly i.v. administration (n=3 animals per group) at 0.3 and 1.0 mg/kg dose. **e-j,** Corresponding changes in total reticulocyte count, aspartate aminotransferase (AST), neutrophils, complement and cytokine activation profiles. Data is presented as mean ± stdev.

To evaluate repeated dosing, monkeys received weekly injections of hPep/hEPO PNPs at 0.3 mg/kg and 1 mg/kg for three weeks. hEPO expression kinetics were measured at 6-, 24-, 72-, and 168-hours post-injection. As shown in Fig. 6d, hPep3 PNPs maintained consistent pharmacodynamics across all doses and injections. Peak hEPO levels were sustained without loss of efficacy, confirming that repeated dosing did not trigger neutralizing responses or toxicity that would limit protein expression. This consistency is crucial for applications requiring sustained protein expression. Reticulocyte counts increased at 72 h and 168 h post-dosing (Fig. 6e and Supplementary Fig. 12a), while RBCs, hemoglobin, and hematocrit remained stable (Supplementary Fig. 12b-d).

Safety assessments confirmed overall tolerability. No clinical signs of toxicity or changes in body weight were observed at doses up to 1 mg/kg in either single or repeated dosing (data not shown). At 1 mg/kg, only mild and transient changes were noted, including increases in aspartate aminotransferase (AST), neutrophils (Fig. 6f,g), prothrombin time (PT), activated partial thromboplastin time (APTT) (Supplementary Fig. 12e,f), complement factors Bb and C3a fragments, cytokine IL-6 (Fig. 6h-j), along with slight decrease in lymphocytes and eosinophils (Supplementary Fig. 12g,h). IFN-γ and TNF-α levels, like alanine transaminase (ALT) and alkaline phosphatase (ALP) were unaffected (data not shown and Supplementary Fig. 12i,j). Control formulation with non-hEPO coding mRNA produced similar responses during the 24-hour observation window (Supplementary Fig. 13a,b). Based on these findings, the No Adverse Effect Level (NOAEL) was established at 1 mg/kg/day.

Together, these studies underscore the potential of hPep3 as a safe and efficient vehicle for mRNA delivery in higher biological systems. Its ability to induce dose-dependent protein expression, maintain efficacy during repeated dosing, and elicit only mild, transient laboratory changes highlights its potential for therapeutic development. Future work will focus on optimizing dosing regimens, assessing long-term safety, and evaluating efficacy in disease-specific models.

## Discussion

The therapeutic use of mRNA without a carrier system is hindered by its poor bioavailability, rapid degradation (with a half-life in minutes), and immunogenicity [34]. To address these challenges, delivery systems such as LNPs have been developed, currently serving as the standard for mRNA delivery. Despite their high efficacy, LNPs predominantly target the liver, even when administered through various routes, limiting their application for extra-targeting [12]. Furthermore, mRNA LNPs often induce a mild to moderate proinflammatory response, an effect linked to the LNP composition and seen in both siRNA and mRNA LNP studies in preclinical and clinical settings [35]. Alternative delivery systems that can minimize these side effects would be particularly advantageous for chronic diseases requiring repeated dosing. Here, we demonstrate that peptide-based nanoparticles, specifically those formulated with hPep3, effectively encapsulate and deliver mRNA while exhibiting a favorable safety profile even after repeated administration.

hPep peptides are designed as alpha-helical, amphipathic peptides, with positive charges positioned on one side of the helix and hydrophobic residues on the other. This amphipathicity was enhanced by incorporating hydrophobic amino acid analogs, such as alkenyl-alanines, into the hydrophobic side of the helix, which increased the overall hydrophobicity. Our in vitro and in vivo screening studies revealed that only certain alkenyl-analogs, particularly octenyl-alanines, provided significant efficacy benefits, while others, like fatty acid-modified lysines, were less effective despite enabling efficient mRNA encapsulation into nanoparticles. Validation studies with hPep3, which contains three octenyl-alanines, demonstrated a high tolerance in mice and effective mRNA transfection both locally and systemically. While some other analogs, like hPep3-OctG and hPep3-2Ado showed promising results in vivo, hPep3 consistently demonstrated the highest efficacy. Mechanistic studies on the cellular uptake of hPep3/mRNA PNPs showed fast energy-dependent endocytic uptake via several pathways. As confirmed by colocalization studies, the hPep3 PNPs localized in lysosomes within a few hours, while endosomal release was mainly detected after 4-24 hours. Together, these promising results supported the progression to NHP studies.

To scale up hPep3 PNPs formulation for NHP studies, we utilized microfluidic mixing to achieve a consistent, sub-100 nm nanoparticle size with two different mRNAs, hEPO and FLuc (859 and 1922 bases, respectively). Downstream processing involved purification and concentration using TFF and sterile filtration, and dilution into cryostorage buffer. The hPep3 PNPs consistently exhibited spherical shape, narrow size distribution (PDIs around 0.170), positive zeta potential, and encapsulation efficiency exceeding 90%. Overall production yields were over 70% in terms of mRNA recovery, which is considerably higher than typically reported for LNPs [36]. The formulation maintained its properties after cryostorage, confirming that this scalable method is suitable for producing peptide-based formulations. Compared to LNPs, the PNPs also offer advantages in formulation simplicity, as only one component is required for mRNA encapsulation and delivery instead of four (ionizable lipid, helper lipid, cholesterol, and PEG-lipid), and no organic solvents are needed [37], which drastically simplifies the process. Validation of microfluidically formulated hPep3 PNPs with FLuc mRNA demonstrated equal or greater luciferase expression in mice than the bulk mixed PNPs, marking an improvement over the parent formulation.

Safety and pharmacokinetic studies in NHPs demonstrated that hPep3 PNPs were well tolerated and effective. Both 0.3 mg/kg and 1 mg/kg doses were administered by single or repeated weekly injections over three weeks, without clinical signs of toxicity. At 1 mg/kg dosing, only mild transient changes were observed, including increases in neutrophils, coagulation parameters (PT, aPTT), liver enzyme (AST), together with modest elevations in complement factors C3a, Bb, and cytokine IL-6. IFN-γ and TNF-α remained unaffected, and all changes resolved spontaneously. Importantly, none of these changes were detected at 0.3 mg/kg, supporting a favorable safety profile even with repeated dosing. Compared to LNPs, which often require extended infusion times and still induce strong immune activation [38–41], hPep3 PNPs appear less reactogenic and more practical for systemic administration.

Pharmacokinetic analyses further confirmed functional delivery. hPep3/hEPO PNPs induced clear dose-dependent increase in serum EPO production, while sustaining consistent levels after repeated dosing, indicating effective long-term protein expression without significant adverse effects, crucial for therapeutic applications. Based on literature, for hPep3 PNPs at 1 mg/kg the peak serum hEPO concentrations were comparable to those achieved with MC3 LNPs at 0.01 mg/kg [38,39] and C12-200 LNPs at 0.05 mg/kg [42] and 2-fold lower than Lipid 1 LNPs at 0.1 mg/kg [40], making them roughly 100-fold, 20-fold and 20-fold more efficient at peak protein induction, respectively. Expression kinetics paralleled those of LNPs, with peak expression at around 6 h, suggesting that the mRNA cargo primarily determines the kinetics rather than the carrier composition.

Biodistribution studies using Cy5-labelled mRNA revealed a clear divergence between nanoparticle localization and transgene expression. In the lungs, only about 8% of hPep3 PNPs were localized, yet they accounted for roughly 70% of total luciferase output, indicating highly efficient translation in pulmonary tissue. Conversely, about 60% of PNPs accumulated in the liver but contributed only about 2% of expression, suggesting tissue-specific differences in mRNA stability or translation efficiency. In the spleen, localization and expression were proportional at around 20%. Compared to LNPs, which show strong hepatocyte tropism and limited extrahepatic expression [11,12], hPep3 PNPs achieve broader distribution, with particularly strong expression in lungs and spleen. This versatility highlights the complementary nature of the two platforms: LNPs excel in hepatocyte-targeted protein production, whereas PNPs may be more suited for multi-organ or immune-modulating applications. Despite the exemplary efficacy of LNPs in soluble protein production, PNPs offer complementary strengths, and ongoing peptide engineering is likely to close the performance gap, paralleling the leap in efficacy achieved with ionizable lipids [43].

Protein corona analysis revealed interactions between hPep3/mRNA PNPs and specific plasma proteins, with notable enrichment of APOE and VTN. These proteins may influence tropism toward the lungs and liver [44]. Understanding and optimizing these interactions will be essential for enhancing therapeutic targeting and efficacy while minimizing off-target effects.

## Conclusions and Future Directions

In conclusion, hPep3-based PNPs present a versatile and promising alternative to traditional lipid-based mRNA delivery systems. Designed to overcome key limitations of RNA delivery, hPep3 PNPs have demonstrated efficient mRNA encapsulation, favorable biodistribution, and targeted delivery with a minimal proinflammatory response. The safety and efficacy profiles observed in NHP studies underscore their potential for broader therapeutic applications, especially in cases requiring multi-organ targeting or repeated dosing in chronic conditions. Moreover, the selective targeting of immune cells and the demonstrated CNS delivery capabilities highlight hPep3 PNPs as a platform with substantial utility in immune modulation and neurological therapies.

Future research will focus on optimizing hPep3 PNPs for enhanced tissue specificity and therapeutic efficacy. Key directions include refining peptide design to achieve targeted delivery to specific cell types within organs and further improving delivery efficiency to match or exceed that of LNPs. In particular, modifying the hydrophobic and cationic domains of hPep3 could improve cellular uptake and endosomal escape, potentially leading to increased transfection efficacy in diverse tissue types. Additionally, exploring the role of the protein corona and optimizing NP interactions with plasma proteins such as APOE and VTN could further enhance targeting precision and minimize off-target effects.

With ongoing advancements in peptide engineering and microfluidic formulation techniques, hPep3 PNPs are well-positioned to advance into clinical development. Upcoming studies will aim to evaluate long-term safety and efficacy in disease-specific models, particularly for applications in immune-related diseases and CNS disorders. By building on these initial findings, hPep3 PNPs have the potential to establish themselves as a safe and efficient platform for mRNA therapeutics, addressing current limitations and expanding the clinical reach of mRNA-based therapies.

## Methods

1. Materials

All peptides were ordered from Pepscan Presto (The Netherlands) with >90% purity. The peptides are C-terminally amidated and have a free amine in the N-terminus. The purities of the peptides were obtained by UPLC (Ultra Performance Liquid Chromatography). Chemically modified Firefly Luciferase (FLuc), Enhanced Green Fluorescent Protein (EGFP) were purchased from TriLink Biotechnologies (USA), Cy5-FLuc mRNA and mCherry mRNA were purchased from APExBIO Technology LLC (USA). All mRNAs were fully substituted with 5-methoxyuridine and modified with CleanCap®.

2. Formulation of peptide/mRNA PNPs

Peptide/mRNA PNPs were formulated in MQ water or 10 mM HEPES buffer at different peptide/mRNA charge ratios (CRs) 1:1-4:1. After adding the peptide to the mRNA the PNPs were let to incubate at RT for 30 min. As a positive control, Lipofectamine^TM^2000 (Invitrogen, Sweden) was used according to manufacturer’s protocol.

3. Encapsulation efficiency and complex stability studies

mRNA condensation and incorporation into PNPs was analyzed by RiboGreen (Thermo Scientific, USA) exclusion assay. Briefly, PNPs were formed as described above. 50 μl of PNPs, formulated at different CRs, were transferred into a black 96-well plate (NUNC, Sweden) containing 140 μl of MQ water and 10 μl of RiboGreen solution. The RiboGreen was let to intercalate for 15 min followed by fluorometric analysis on a SynergyMx (BioTek, USA) at 485 nm excitation and 528 nm emission wavelengths. Results are normalized to the fluorescence of naked mRNA.

For the analysis of peptide/mRNA PNP resistance to competitive anion heparin treatment, preformed peptide/mRNA PNPs containing 0.1 µg of mRNA were incubated in the presence of heparin (Sigma-Aldrich, Germany) and RiboGreen. The fluorescence of RiboGreen bound to exposed mRNA was measured over time. Results are expressed as relative fluorescence, where 100% is the fluorescence of naked mRNA treated with heparin. The measurements were carried out in duplicates.

4. DLS

Z-average size, polydispersity index (PDI) and zeta potential of peptide/mRNA PNPs were determined by dynamic light scattering using a Zetasizer Ultra Red apparatus (Malvern Panalytical, United Kingdom). Peptide/mRNA PNPs were formulated as described above and measured in ZEN0040 disposable cuvettes (Malvern Panalytical). Each measurement was repeated three times in a row while one measurement was set to 8 runs (1 run was 8 s). Prior to zeta potential measurement PNPs were diluted to 800 μl with 10 mM HEPES buffer pH7.4 and transferred to DTS1070 folded capillary zeta cell cuvettes (Malvern Panalytical). The measurement parameters were set to automatic. All measurements were carried out at 22°C.

5. Cell cultures and transfection

Cells were cultured in high glucose Dulbecco’s modified Eagle’s medium (DMEM) (Gibco®, ThermoFisher) supplemented with GlutaMAX^TM^, sodium pyruvate plus 10% fetal bovine serum, and 1% penicillin-streptomycin (Gibco®, ThermoFisher) in a water-jacketed incubator at 37 °C and 5% CO2. For the transfection efficiency experiments 10,000 cells per well were seeded 24 h prior to transfection into transparent, tissue culture-treated, clear flat-bottom 96-well plates (Corning) in final volume of 100 µl. On the day of transfection, 10 µl of preformed hPep/mRNA PNPs at 100 ng final mRNA dose were added to cells in triplicates and treated for 24 h. After the treatment, media was removed and cells were lysed in 0.1% Triton X-100 (Sigma-Aldrich) for 30 min at room temperature. The luciferase activity was measured from the lysates with the Luciferase Assay Kit (Promega, Madison, WI, USA) under GLOMAX 96 microplate luminometer (Promega) and normalized to the protein content (Bio-Rad Protein Assay Kit II, BioRad, Hercules, CA, USA).

6. Flow cytometry analysis

20,000 cells per well were seeded on 96-well plates 24 h prior to treatment. For dose titration studies, the cells were treated with Cy5-mRNA or hPep/Cy5-mRNA PNPs at CR4 at the dose range of 3.125-100 ng/well in serum-containing conditions for 4 h. For uptake experiments, hPep/Cy5-mRNA PNPs (CR4) at 50 ng/well of mRNA were used to treat the cells for 15 min, 1 hour, or 4 hours in serum-containing conditions.

For mCherry expression analysis cells were treated with hPep3/EZ Cap™ mCherry mRNA (ApexBio, USA) NPs (CR4) at 200 ng/well for 24 hours. After treatment, the samples were prepared for flow cytometry analysis using standard preparation procedures. For live/dead cell exclusion the cells were stained with 0.5 µg/ml DAPI solution (Thermo Fisher Scientific) and measured on Attune NxT Flow Cytometer (Thermo Fisher Scientific). The data was analyzed using FlowJo Software 10.8.1 (BD Biosciences, USA).

For endocytosis inhibition studies 20,000 cells per well were seeded on 96-well plates 24 h prior to treatment. Next day cells were pre-incubated with endocytosis inhibitors for 1h: a mixture of 5 mM of sodium azide and 3 mM D-glucose (-ATP) for energy-dependent endocytosis inhibition, 25 µM nystatin (Nys) for caveolae-mediated endocytosis, 15 µM chlorpromazine (CPZ) for clathrin-mediated endocytosis and 0.25 µM cytochalasin D (Cyt D) for macropinocytosis. Thereafter, cells were treated with hPep/Cy5-mRNA PNPs (CR4) at 50 ng/well of mRNA for 3 hours, followed by sample preparation as described above. Data were normalized to uptake without inhibitors (hPep3) and are presented as percentage of initial uptake.

7. Microscopy

For uptake experiments, A549 cells (50,000 per well) were seeded onto round glass coverslips (VWR, Germany) in 24-well cell culture plates. After 48 hours, cells were treated with 250 ng Cy5-mRNA alone or with peptide/Cy5-mRNA PNPs at CR4 for 15 min to 4 hours at 37 ℃. Then, cells were washed twice with PBS and fixed in 4% paraformaldehyde in 0.1 M PBS (pH7.4) for 15 min at room temperature. Fixed cells were washed twice with PBS, and nuclei were stained with Hoechst 33342 solution (1 µg/ml in PBS; Thermo Fischer Scientific) for 5 min at room temperature. Coverslips were mounted on Superfrost Plus slides (Epredia, USA) and imaged using Zeiss LSM900 confocal microscope (Carl Zeiss, Germany).

For colocalization studies, A549 cells (5,000 per well) were seeded into 18-well chambers (Ibidi, Germany) 48 hours before the treatment. After incubation with peptide/Cy5-mRNA PNPs (CR4) at 45 ng/well of mRNA, 50 nM LysoTracker-Green (Thermo Fischer Scientific) was added, and the cells were visualized in 5% CO_2_ chamber at 37 °C.

For mCherry expression, A549 cells (30,000 per well) were seeded into 8-well chambers (Ibidi, Germany) 48 hours before the treatment. After incubation the cells were treated with hPep3/EZ Cap™ mCherry mRNA (ApexBio, USA) NPs (CR4) at 625 ng/well for 4 hours in serum-free media, followed by 20 hour-incubation in serum-containing conditions.

For endosomal escape, HeLa-galectin-8 (Gal8)-mRuby3 cell line, stably expressing mRuby3-Gal8 fusion protein, was created [45,46] and 50,000 cells per well were seeded onto round glass coverslips in 24-well cell culture plates 48 hours before the experiment. After the incubation with peptide/Cy5-mRNA PNPs at 250 ng/well of mRNA, the cells were fixed, and nuclei were stained as described above.

8. Animal studies

The in vivo experiments in mice were approved by the Swedish local laboratory animal research ethics committee (approval no. S4-16) and the Estonian Laboratory Animal Ethics Committee (approval no. 185 and 1.2-13/19), and all experiments were performed in accordance with relevant guidelines and regulations. To measure mRNA delivery efficacy in vivo, female NMRI or Balb/c mice with a bodyweight of around 20 g were IV (tail vein) or ICV (as described previously) injected with modified mRNA encoding for firefly luciferase (Trilink Biotechnologies, USA) and formulated either with CPPs or LNPs (as previously described). For assessing luciferase activity in vivo, animals were imaged by IVIS Spectrum (Perkin Elmer, USA) for firefly luciferase luminescence after IP administration of 150 mg/kg D luciferin (Perkin Elmer, USA). Here, either live (isoflurane sedated) mice were imaged, or the animals were sacrificed, and the organs harvested prior to analysis. All in vivo data were processed by IVIS software (Living Image Software, Perkin Elmer, USA). Adobe Photoshop CS4 and Adobe Illustrator (Adobe, USA) were used to crop out and align the organ images.

For assessing biodistribution and bioavailability of hPep/mRNA PNPs, female NMRI mice with a bodyweight of around 20 g were IV (tail vein) injected with Cy5-labelled modified mRNA encoding for firefly luciferase (Trilink Biotechnologies, USA) formulated with CPPs. For assessing luciferase activity in vivo, animals were imaged by IVIS Spectrum (Perkin Elmer, USA) for firefly luciferase luminescence after IP administration of 150 mg/kg D-luciferin. Here, either live (isoflurane sedated) mice were imaged, or the animals were sacrificed, and the organs harvested prior to analysis. Whereas for biodistribution, Cy5 fluorescent signal from harvested organs was quantified using IVIS.

9. FACS for cell typing

To determine the target cell type of hPep/mRNA PNPs, modified mRNA encoding EGFP was used. Followed by intravenous administration of PNPs in NMRI female mice (about 20 g), after 24 hours, animals were euthanized, and different organs were harvested. The lung and spleen were dissociated into single-cell suspension by mechanical disruption, and isolated cells were subjected to flow cytometry.

10. NHP handling and treatment

Nonhuman primate studies were conducted at ITR Laboratories Canada Inc. (Baie d’Urfe, Quebec, Canada) using female cynomolgus monkeys (Macaca Fascicularis), young adults weighing 2.0–4.0 kg.

Animals were housed in groups of up to 3, in stainless steel, perforated-floor cages, in a temperature-and humidity-controlled environment (21°C–26°C and 30%–70%, respectively), with an automatic 12-h dark/light cycle. Purified tap water and food (LabDiet Certified Primate Diet #5048) were available to animals *ad libitum*. Concentrations of the dietary constituents and environmental contaminants were routinely monitored. Tuberculin tests were carried out upon arrival at the test facility. Animals were given access to toys throughout the study, except during designated procedures.

The study plan and relevant amendments for this study were reviewed and assessed by the Animal Care Committee (ACC) of ITR. All animals used in this study were cared for in accordance with the principles outlined in the current “Guide to the Care and Use of Experimental Animals” as published by the Canadian Council on Animal Care and the “Guide for the Care and Use of Laboratory Animals”, an NIH publication. No treatment randomization or blinding methods were used for any of the animal studies. Sample sizes were determined by the resource equation method (n=3 per group).

hPep/mRNA NPs were administered in HEPES-buffered sucrose (pH7.4) by slow bolus IV injection over a period of 60 to 90 seconds via a disposable indwelling catheter placed in the saphenous using a hypodermic needle attached to a syringe. The dose volume was 5 mL/kg for all animals.

11. Blood collection from NHPs

Blood samples were collected by venipuncture (other than the one used for dosing) following an overnight period of food deprivation. Various clinical pathology parameters that are automatically recorded by the instrumentation were not reported. Hematology parameters were measured from approximately 0.3 mL blood samples collected into K2EDTA anticoagulant containing tubes. Coagulation parameters activated partial thromboplastin time and prothrombin time were measured from approximately 0.75 mL blood samples collected into sodium citrate anticoagulant. Clinical chemistry parameters were measured from approximately 0.75 mL blood samples collected into tubes containing a clotting activator.

12. Complement analysis (C3a, Bb)

Blood samples (approximately 0.8 mL) were collected from each monkey during the pre-treatment period. For this purpose, each monkey was bled by venipuncture (other than the one used for dosing), and the samples were collected into tubes containing the anticoagulant, K_2_EDTA. The K_2_EDTA tubes were placed on wet ice pending processing. Bb fragment concentration was quantified in monkey plasma using an enzyme-linked immunosorbent assay (ELISA), MicroVue^TM^ Complement Bb Plus EIA Kit (Cat. #: A027, QuidelOrtho Corporation, USA) at ITR. C3a fragment concentration was quantified in monkey plasma using a MicroVue^TM^ Complement C3a Plus EIA Kit (Cat. #: A031, QuidelOrtho Corporation, USA) at ITR. Raw data was acquired using the Powerwave XS plate reader with the Biotek Gen5™ Secure software Version 3.03.14 (Agilent, USA).

13. Cytokine analysis (TNF α, IL-6 and IFN-γ)

Blood samples (approximately 0.3 mL each) were collected as described above but the samples were collected into tubes containing clotting activator. Serum samples were analyzed for TNF α, IL-6 and IFN-γ concentrations at ITR using an electrochemiluminescent multiplex immunoassay (Meso Scale Diagnostics, USA) according to the manufacturer’s instructions. The electrochemiluminescence readings were acquired with the Meso Scale Discovery QuickPlex SQ 120 instrument operated with the Meso Scale Discovery WORKBENCH software version 4 (Meso Scale Diagnostics LLC, USA).

14. hEPO analysis

Blood samples (approximately 0.3 mL each) were collected as described above but the samples were collected into tubes containing no anticoagulant. Following collection, blood samples were allowed to stand at room temperature for approximately 30 minutes to clot. The samples were then centrifuged (2,500 rpm for 10 minutes at approximately 4 °C) and the resulting serum was recovered, divided into two aliquots (sets 1 and 2), and stored frozen (≤ - 60 °C) in appropriately labeled tubes pending transfer for analysis. The serum samples were analyzed at ITR using an Erythropoietin (EPO) ELISA kit (Cat. #:01630, STEMCELL Technologies Inc., Canada).

15. Statistical analysis

CLSM images were analyzed using Fiji [47], for colocalization BIOP JACoP plugin was used [48]. Data are presented as mean with a standard error of the (mean ± SEM) of at least three independent experiments. Significant differences were evaluated by analysis of variance (ANOVA) with Dunnett’s multiple comparison test, Bonferronís multiple comparison test, Kruskal-Wallis test, or unpaired t test (GraphPad Prism 8 or 9; GraphPad Software, Inc., USA). In all cases, differences with p < 0.05 were deemed to be significant (* p < 0.05, ** p < 0.01, *** p < 0.001, and **** p < 0.0001).

## Acknowledgements

T.L. (Taavi Lehto) was supported by Estonian Research Council grants PSG226 and PRG1882. S.E.A. was supported by the H2020 (EXPERT); Swedish foundation of Strategic Research FormulaEx, SM19-0007; European Research Council (ERC) under the European Union’s Horizon 2020 research and innovation programme (DELIVER, grant agreement No 101001374); Cancerfonden; Swedish Research Council grant 4–258/2021. T.L. and S.E.A. acknowledge the support from Evox Therapeutics. Helena Sork discloses support from the Estonian Research Council grant PSG1043. Biorender.com has been used to create visuals for the Figures.

## References

1. Sahin, U.; Karikó, K.; Türeci, Ö. MRNA-Based Therapeutics-Developing a New Class of Drugs. Nat. Rev. Drug Discov. 2014, 13, 759–780.

2. Miao, L.; Zhang, Y.; Huang, L. MRNA Vaccine for Cancer Immunotherapy. Mol. Cancer 2021, 20, doi:10.1186/s12943-021-01335-5.

3. Van Hoecke, L.; Verbeke, R.; Dewitte, H.; Lentacker, I.; Vermaelen, K.; Breckpot, K.; Van Lint, S. MRNA in Cancer Immunotherapy: Beyond a Source of Antigen. Mol. Cancer 2021, 20, doi:10.1186/s12943-021-01329-3.

4. Hunter, T.L.; Bao, Y.; Zhang, Y.; Matsuda, D.; Riener, R.; Wang, A.; Li, J.J.; Soldevila, F.; Chu, D.S.H.; Nguyen, D.P.;, et al. In Vivo CAR T Cell Generation to Treat Cancer and Autoimmune Disease. Science (80-.). 2025, 388, 1311–1317, doi:10.1126/science.ads8473.

5. Sahu, I.; Haque, A.K.M.A.; Weidensee, B.; Weinmann, P.; Kormann, M.S.D. Recent Developments in MRNA-Based Protein Supplementation Therapy to Target Lung Diseases. Mol. Ther. 2019, 27, 803–823, doi:10.1016/j.ymthe.2019.02.019.

6. Barbier, A.J.; Jiang, A.Y.; Zhang, P.; Wooster, R.; Anderson, D.G. The Clinical Progress of MRNA Vaccines and Immunotherapies. Nat. Biotechnol. 2022, 40, 840–854, doi:10.1038/s41587-022-01294-2.

7. Li, B.; Manan, R.S.; Liang, S.Q.; Gordon, A.; Jiang, A.; Varley, A.; Gao, G.; Langer, R.; Xue, W.; Anderson, D. Combinatorial Design of Nanoparticles for Pulmonary MRNA Delivery and Genome Editing. Nat. Biotechnol. 2023, 41, 1410–1415, doi:10.1038/s41587-023-01679-x.

8. Rowe, S.M.; Zuckerman, J.B.; Dorgan, D.; Lascano, J.; McCoy, K.; Jain, M.; Schechter, M.S.; Lommatzsch, S.; Indihar, V.; Lechtzin, N.;, et al. Inhaled MRNA Therapy for Treatment of Cystic Fibrosis: Interim Results of a Randomized, Double-blind, Placebo-controlled Phase 1/2 Clinical Study. J. Cyst. Fibros. 2023, 22, 656–664, doi:10.1016/j.jcf.2023.04.008.

9. Koeberl, D.; Schulze, A.; Sondheimer, N.; Lipshutz, G.S.; Geberhiwot, T.; Li, L.; Saini, R.; Luo, J.; Sikirica, V.; Jin, L.;, et al. Interim Analyses of a First-in-Human Phase 1/2 MRNA Trial for Propionic Acidaemia. Nature 2024, 628, 872–877, doi:10.1038/s41586-024-07266-7.

10. Hou, X.; Zaks, T.; Langer, R.; Dong, Y. Lipid Nanoparticles for MRNA Delivery. Nat. Rev. Mater. 2021, 6, 1078–1094.

11. Zhang, Y.N.; Poon, W.; Tavares, A.J.; McGilvray, I.D.; Chan, W.C.W. Nanoparticle–Liver Interactions: Cellular Uptake and Hepatobiliary Elimination. J. Control. Release 2016, 240, 332–348, doi:10.1016/j.jconrel.2016.01.020.

12. Loughrey, D.; Dahlman, J.E. Non-Liver MRNA Delivery. Acc. Chem. Res. 2022, 55, 13–23, doi:10.1021/acs.accounts.1c00601.

13. Peng, L.; Wagner, E. Polymeric Carriers for Nucleic Acid Delivery: Current Designs and Future Directions. Biomacromolecules 2019, 20, 3613–3626, doi:10.1021/acs.biomac.9b00999.

14. Berger, S.; Lächelt, U.; Wagner, E. Dynamic Carriers for Therapeutic RNA Delivery. Proc. Natl. Acad. Sci. U. S. A. 2024, 121, e2307799120, doi:10.1073/pnas.2307799120.

15. Lehto, T.; Ezzat, K.; Wood, M.J.A.; EL Andaloussi, S. Peptides for Nucleic Acid Delivery. Adv. Drug Deliv. Rev. 2016, 106, 172–182, doi:10.1016/j.addr.2016.06.008.

16. Langel, Ü. Cell-Penetrating Peptides: Methods and Protocols; Langel, Ü., Ed.; Methods in Molecular Biology; Springer New York: New York, NY, 2015; Vol. 1324; ISBN 9781493928064.

17. Betts, C.; Saleh, A.F.; Arzumanov, A.A.; Hammond, S.M.; Godfrey, C.; Coursindel, T.; Gait, M.J.; Wood, M.J. Pip6-PMO, a New Generation of Peptide-Oligonucleotide Conjugates with Improved Cardiac Exon Skipping Activity for DMD Treatment. Mol. Ther. - Nucleic Acids 2012, 1, e38, doi:10.1038/mtna.2012.30.

18. EL Andaloussi, S.; Lehto, T.; Mäger, I.; Rosenthal-Aizman, K.; Oprea, I.I.; Simonson, O.E.; Sork, H.; Ezzat, K.; Copolovici, D.M.; Kurrikoff, K.;, et al. Design of a Peptide-Based Vector, PepFect6, for Efficient Delivery of SiRNA in Cell Culture and Systemically in Vivo. Nucleic Acids Res. 2011, 39, 3972–3987, doi:10.1093/nar/gkq1299.

19. Porosk, L.; Härk, H.H.; Arukuusk, P.; Haljasorg, U.; Peterson, P.; Kurrikoff, K. The Development of Cell-Penetrating Peptides for Efficient and Selective In Vivo Expression of MRNA in Spleen Tissue. Pharmaceutics 2023, 15, 952, doi:10.3390/pharmaceutics15030952.

20. Bestas, B.; Moreno, P.M.D.; Blomberg, K.E.M.; Mohammad, D.K.; Saleh, A.F.; Sutlu, T.; Nordin, J.Z.; Guterstam, P.; Gustafsson, M.O.; Kharazi, S.;, et al. Splice-Correcting Oligonucleotides Restore BTK Function in X-Linked Agammaglobulinemia Model. J. Clin. Invest. 2014, 124, 4067–4081, doi:10.1172/JCI76175.

21. Gros, E.; Deshayes, S.; Morris, M.C.; Aldrian-Herrada, G.; Depollier, J.; Heitz, F.; Divita, G. A Non-Covalent Peptide-Based Strategy for Protein and Peptide Nucleic Acid Transduction. Biochim. Biophys. Acta - Biomembr. 2006, 1758, 384–393, doi:10.1016/j.bbamem.2006.02.006.

22. Park, S.E.; Sajid, M.I.; Parang, K.; Tiwari, R.K. Cyclic Cell-Penetrating Peptides as Efficient Intracellular Drug Delivery Tools. Mol. Pharm. 2019, 16, 3727–3743, doi:10.1021/acs.molpharmaceut.9b00633.

23. Futaki, S.; Ohashi, W.; Suzuki, T.; Niwa, M.; Tanaka, S.; Ueda, K.; Harashima, H.; Sugiura, Y. Stearylated Arginine-Rich Peptides: A New Class of Transfection Systems. Bioconjug. Chem. 2001, 12, 1005–1011, doi:10.1021/bc015508l.

24. Lehto, T.; Vasconcelos, L.; Margus, H.; Figueroa, R.; Pooga, M.; Hällbrink, M.; Langel, Ü. Saturated Fatty Acid Analogues of Cell-Penetrating Peptide PepFect14: Role of Fatty Acid Modification in Complexation and Delivery of Splice-Correcting Oligonucleotides. Bioconjug. Chem. 2017, 28, 782–792, doi:10.1021/acs.bioconjchem.6b00680.

25. van Asbeck, A.H.; Dieker, J.; Oude Egberink, R.; van den Berg, L.; van der Vlag, J.; Brock, R. Protein Expression Correlates Linearly with Mrna Dose over up to Five Orders of Magnitude in Vitro and in Vivo. Biomedicines 2021, 9, 511, doi:10.3390/biomedicines9050511.

26. Bazaz, S.; Lehto, T.; Tops, R.; Gissberg, O.; Gupta, D.; Bestas, B.; Bost, J.; Wiklander, O.P.B.; Sork, H.; Zaghloul, E.M.;, et al. Novel Orthogonally Hydrocarbon-Modified Cell-Penetrating Peptide Nanoparticles Mediate Efficient Delivery of Splice-Switching Antisense Oligonucleotides in Vitro and in Vivo. Biomedicines 2021, 9, 1046, doi:10.3390/biomedicines9081046.

27. Lehto, T.; Isakannu, M.; Sork, H.; Lorents, A.; Bazaz, S.; Wiklander, O.P.B.; Andaloussi, S. EL; Lehto, T. Amphipathic Octenyl-Alanine Modified Peptides Mediate Effective SiRNA Delivery. J. Pept. Sci. 2025, 31, e70054, doi:10.1002/PSC.70054.

28. Lehto, T.; Simonson, O.E.; Mäger, I.; Ezzat, K.; Sork, H.; Copolovici, D.M.; Viola, J.R.; Zaghloul, E.M.; Lundin, P.; Moreno, P.M.D.;, et al. A Peptide-Based Vector for Efficient Gene Transfer in Vitro and in Vivo. Mol. Ther. 2011, 19, 1457–1467, doi:10.1038/mt.2011.10.

29. Veiman, K.-L.; Künnapuu, K.; Lehto, T.; Kiisholts, K.; Pärn, K.; Langel, Ü.; Kurrikoff, K. PEG Shielded MMP Sensitive CPPs for Efficient and Tumor Specific Gene Delivery in Vivo. J. Control. Release 2015, 209, 238–247, doi:10.1016/j.jconrel.2015.04.038.

30. Maeki, M.; Kimura, N.; Sato, Y.; Harashima, H.; Tokeshi, M. Advances in Microfluidics for Lipid Nanoparticles and Extracellular Vesicles and Applications in Drug Delivery Systems. Adv. Drug Deliv. Rev. 2018, 128, 84–100.

31. Liu, K.; Salvati, A.; Sabirsh, A. Physiology, Pathology and the Biomolecular Corona: The Confounding Factors in Nanomedicine Design. Nanoscale 2022, 14, 2136–2154, doi:10.1039/d1nr08101b.

32. Liu, R.; Jiang, W.; Walkey, C.D.; Chan, W.C.W.; Cohen, Y. Prediction of Nanoparticles-Cell Association Based on Corona Proteins and Physicochemical Properties. Nanoscale 2015, 7, 9664–9675, doi:10.1039/C5NR01537E.

33. Dilliard, S.A.; Cheng, Q.; Siegwart, D.J. On the Mechanism of Tissue-Specific MRNA Delivery by Selective Organ Targeting Nanoparticles. Proc. Natl. Acad. Sci. U. S. A. 2021, 118, e2109256118, doi:10.1073/pnas.2109256118.

34. Lee, J.; Woodruff, M.C.; Kim, E.H.; Nam, J.H. Knife’s Edge: Balancing Immunogenicity and Reactogenicity in MRNA Vaccines. Exp. Mol. Med. 2023, 55, 1305–1313, doi:10.1038/s12276-023-00999-x.

35. Chaudhary, N.; Kasiewicz, L.N.; Newby, A.N.; Arral, M.L.; Yerneni, S.S.; Melamed, J.R.; LoPresti, S.T.; Fein, K.C.; Strelkova Petersen, D.M.; Kumar, S.;, et al. Amine Headgroups in Ionizable Lipids Drive Immune Responses to Lipid Nanoparticles by Binding to the Receptors TLR4 and CD1d. *Nat*. Biomed. Eng. 2024, 1–16, doi:10.1038/s41551-024-01256-w.

36. Schober, G.B.; Story, S.; Arya, D.P. A Careful Look at Lipid Nanoparticle Characterization: Analysis of Benchmark Formulations for Encapsulation of RNA Cargo Size Gradient. Sci. Rep. 2024, 14, 1–10, doi:10.1038/s41598-024-52685-1.

37. Evers, M.J.W.; Kulkarni, J.A.; van der Meel, R.; Cullis, P.R.; Vader, P.; Schiffelers, R.M. State-of-the-Art Design and Rapid-Mixing Production Techniques of Lipid Nanoparticles for Nucleic Acid Delivery. Small Methods 2018, 1700375, doi:10.1002/smtd.201700375.

38. Sedic, M.; Senn, J.J.; Lynn, A.; Laska, M.; Smith, M.; Platz, S.J.; Bolen, J.; Hoge, S.; Bulychev, A.; Jacquinet, E.;, et al. Safety Evaluation of Lipid Nanoparticle–Formulated Modified MRNA in the Sprague-Dawley Rat and Cynomolgus Monkey. Vet. Pathol. 2018, 55, 341–354, doi:10.1177/0300985817738095.

39. Sabnis, S.; Kumarasinghe, E.S.; Salerno, T.; Mihai, C.; Ketova, T.; Senn, J.J.; Lynn, A.; Bulychev, A.; McFadyen, I.; Chan, J.;, et al. A Novel Amino Lipid Series for MRNA Delivery: Improved Endosomal Escape and Sustained Pharmacology and Safety in Non-Human Primates. Mol. Ther. 2018, 26, 1509–1519, doi:10.1016/j.ymthe.2018.03.010.

40. Cornebise, M.; Narayanan, E.; Xia, Y.; Acosta, E.; Ci, L.; Koch, H.; Milton, J.; Sabnis, S.; Salerno, T.; Benenato, K.E. Discovery of a Novel Amino Lipid That Improves Lipid Nanoparticle Performance through Specific Interactions with MRNA. Adv. Funct. Mater. 2022, 32, doi:10.1002/adfm.202106727.

41. Holland, R.; Lam, K.; Jeng, S.; McClintock, K.; Palmer, L.; Schreiner, P.; Wood, M.; Zhao, W.; Heyes, J. Silicon Ether Ionizable Lipids Enable Potent MRNA Lipid Nanoparticles with Rapid Tissue Clearance. ACS Nano 2024, 18, 10374–10387, doi:10.1021/acsnano.3c09028.

42. DeRosa, F.; Guild, B.; Karve, S.; Smith, L.; Love, K.; Dorkin, J.R.; Kauffman, K.J.; Zhang, J.; Yahalom, B.; Anderson, D.G.;, et al. Therapeutic Efficacy in a Hemophilia B Model Using a Biosynthetic MRNA Liver Depot System. Gene Ther. 2016, 23, 699–707, doi:10.1038/gt.2016.46.

43. Jayaraman, M.; Ansell, S.M.; Mui, B.L.; Tam, Y.K.; Chen, J.; Du, X.; Butler, D.; Eltepu, L.; Matsuda, S.; Narayanannair, J.K.;, et al. Maximizing the Potency of SiRNA Lipid Nanoparticles for Hepatic Gene Silencing In Vivo. Angew. Chemie Int. Ed. 2012, 51, 8529–8533, doi:10.1002/anie.201203263.

44. Dilliard, S.A.; Siegwart, D.J. Passive, Active and Endogenous Organ-Targeted Lipid and Polymer Nanoparticles for Delivery of Genetic Drugs. Nat. Rev. Mater. 2023, 8, 282–300.

45. Rui, Y.; Wilson, D.R.; Tzeng, S.Y.; Yamagata, H.M.; Sudhakar, D.; Conge, M.; Berlinicke, C.A.; Zack, D.J.; Tuesca, A.; Green, J.J. High-Throughput and High-Content Bioassay Enables Tuning of Polyester Nanoparticles for Cellular Uptake, Endosomal Escape, and Systemic in Vivo Delivery of MRNA. Sci. Adv. 2022, 8, 2855, doi:10.1126/sciadv.abk2855.

46. Lin, Y.; Wilk, U.; Pöhmerer, J.; Hörterer, E.; Höhn, M.; Luo, X.; Mai, H.; Wagner, E.; Lächelt, U. Folate Receptor-Mediated Delivery of Cas9 RNP for Enhanced Immune Checkpoint Disruption in Cancer Cells. Small 2023, 19, 2205318, doi:10.1002/smll.202205318.

47. Schindelin, J.; Arganda-Carreras, I.; Frise, E.; Kaynig, V.; Longair, M.; Pietzsch, T.; Preibisch, S.; Rueden, C.; Saalfeld, S.; Schmid, B.;, et al. Fiji: An Open-Source Platform for Biological-Image Analysis. Nat. Methods 2012, 9, 676–682, doi:10.1038/nmeth.2019.

48. Bolte, S.; Cordelières, F.P. A Guided Tour into Subcellular Colocalization Analysis in Light Microscopy. J. Microsc. 2006, 224, 213–232, doi:10.1111/j.1365-2818.2006.01706.x.

