## Supplementary Data for "Peptide nanoparticles for systemic mRNA delivery in rodents and non-human primates"

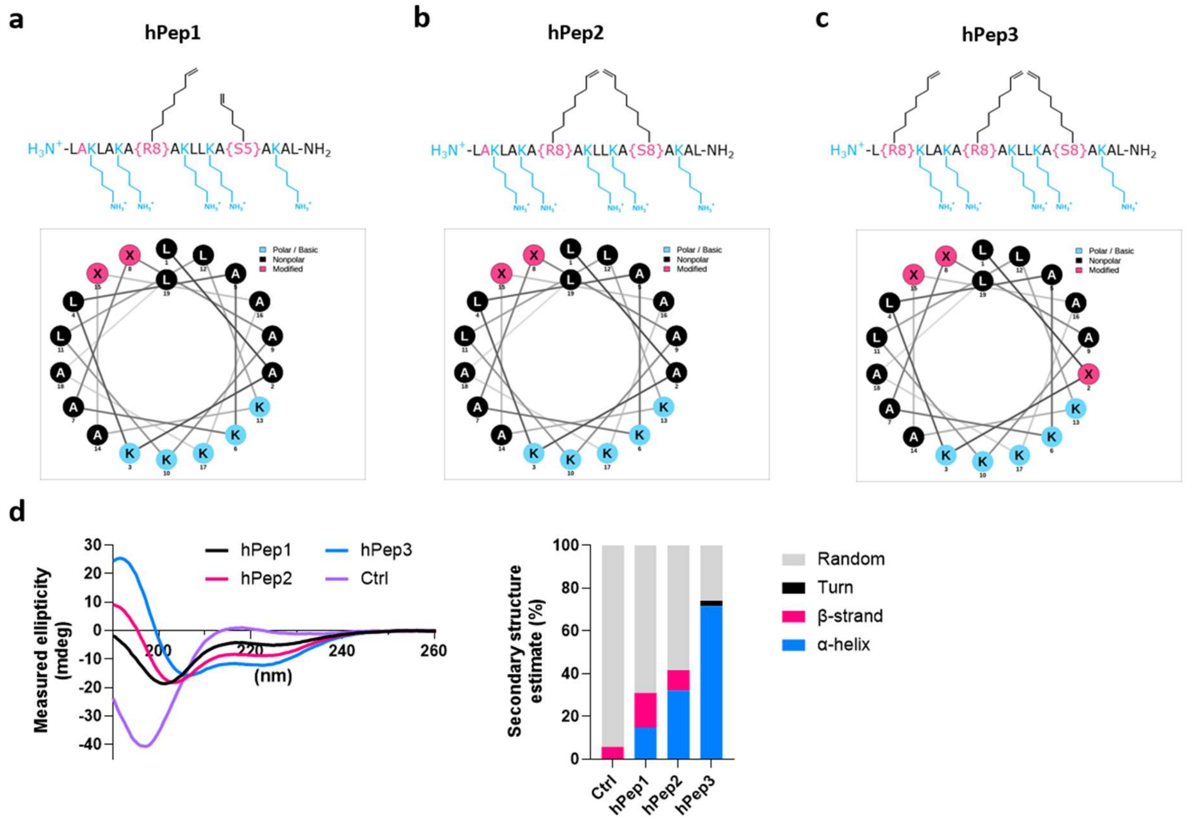


**Supplementary Figure 1. Structural design of hPep peptides. a-c,** Chemical structure and helical wheel projections of hPep peptides. The Ctrl peptide is hPep3 analog where all three octenyl-alanines are substituted by natural alanines. Helical wheel projections were created with http://lbqp.unb.br/NetWheels/. **d,** CD spectra of hPep peptides.

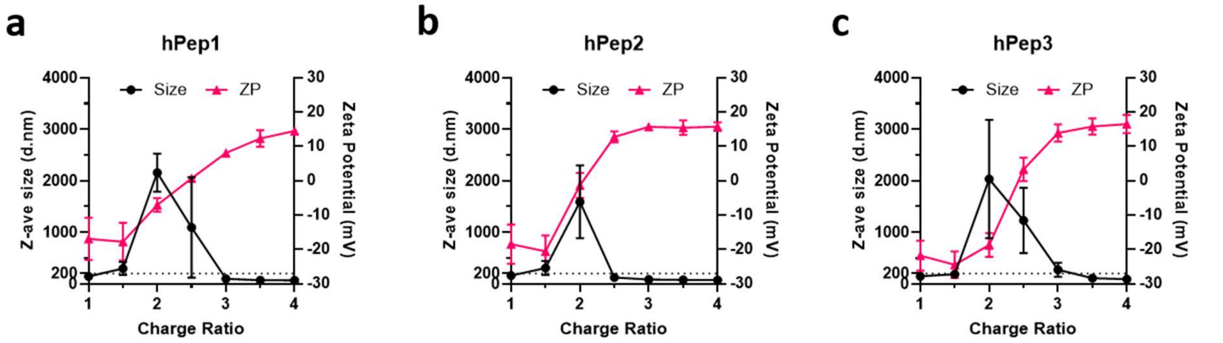


**Supplementary Figure 2. Physicochemical characterization of hPep/mRNA NP properties. a-c,** Size and zeta potential graphs of hPep/mRNA complexes formulated over a range of peptide/mRNA charge ratios with hPep1, hPep2, and hPep3, respectively. Data is presented as mean ± stdev (n=3).

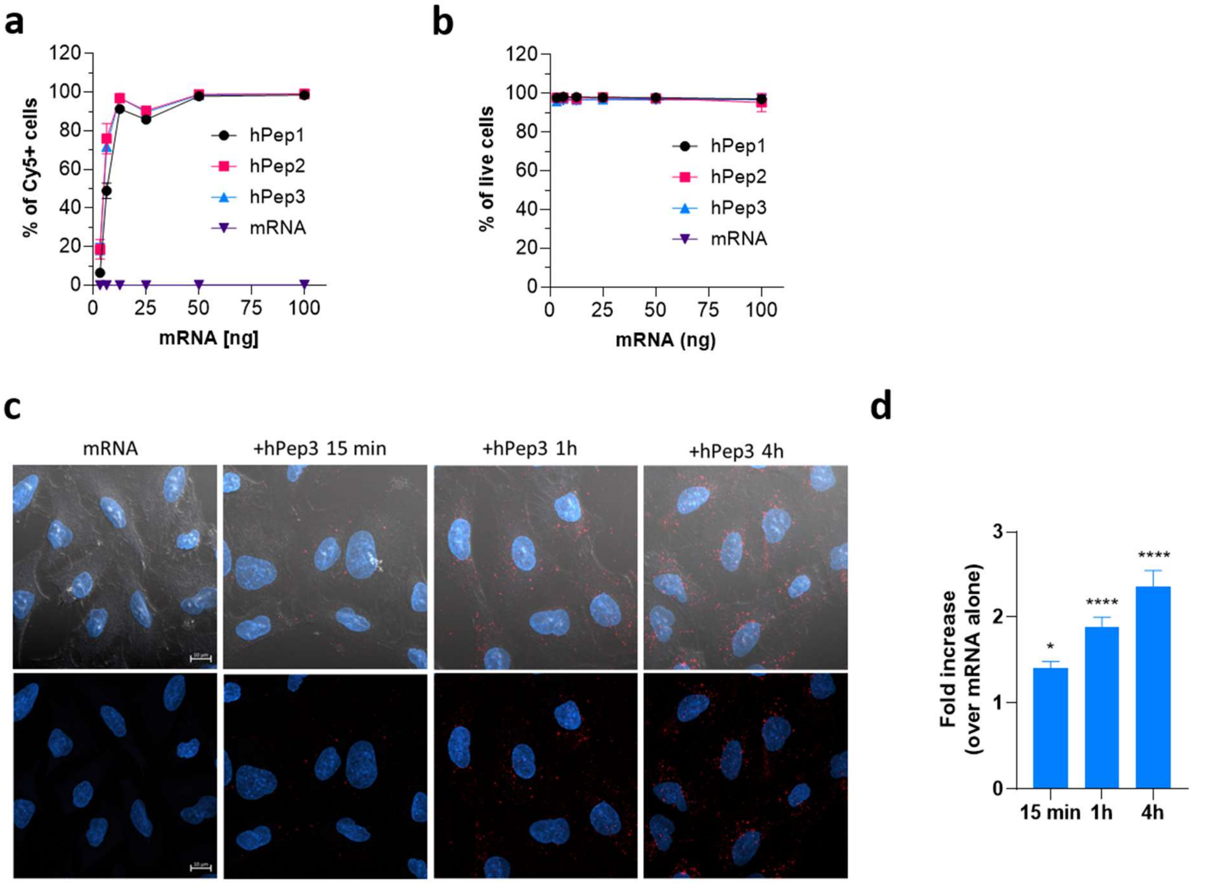


**Supplementary Figure 3. Uptake of hPep/Cy5-mRNA NPs.** A549 cells were treated for 4 h with hPep/Cy5-mRNA NPs formulated at CR4 at different doses. **a**, Percentage of Cy5 positive cells, and **b,** percentage of live cells were quantified by FACS. Data is presented as mean ± stdev (n=3). **c,** Confocal microscopy of fixed A549 cells at different time-points after treatment with hPep3/Cy5-mRNA NPs. **d,** Statistical analysis of the uptake of hPep/Cy5-mRNA NPs (mean ± SEM). Kruskal-Wallis test with Dunn's multiple comparison.

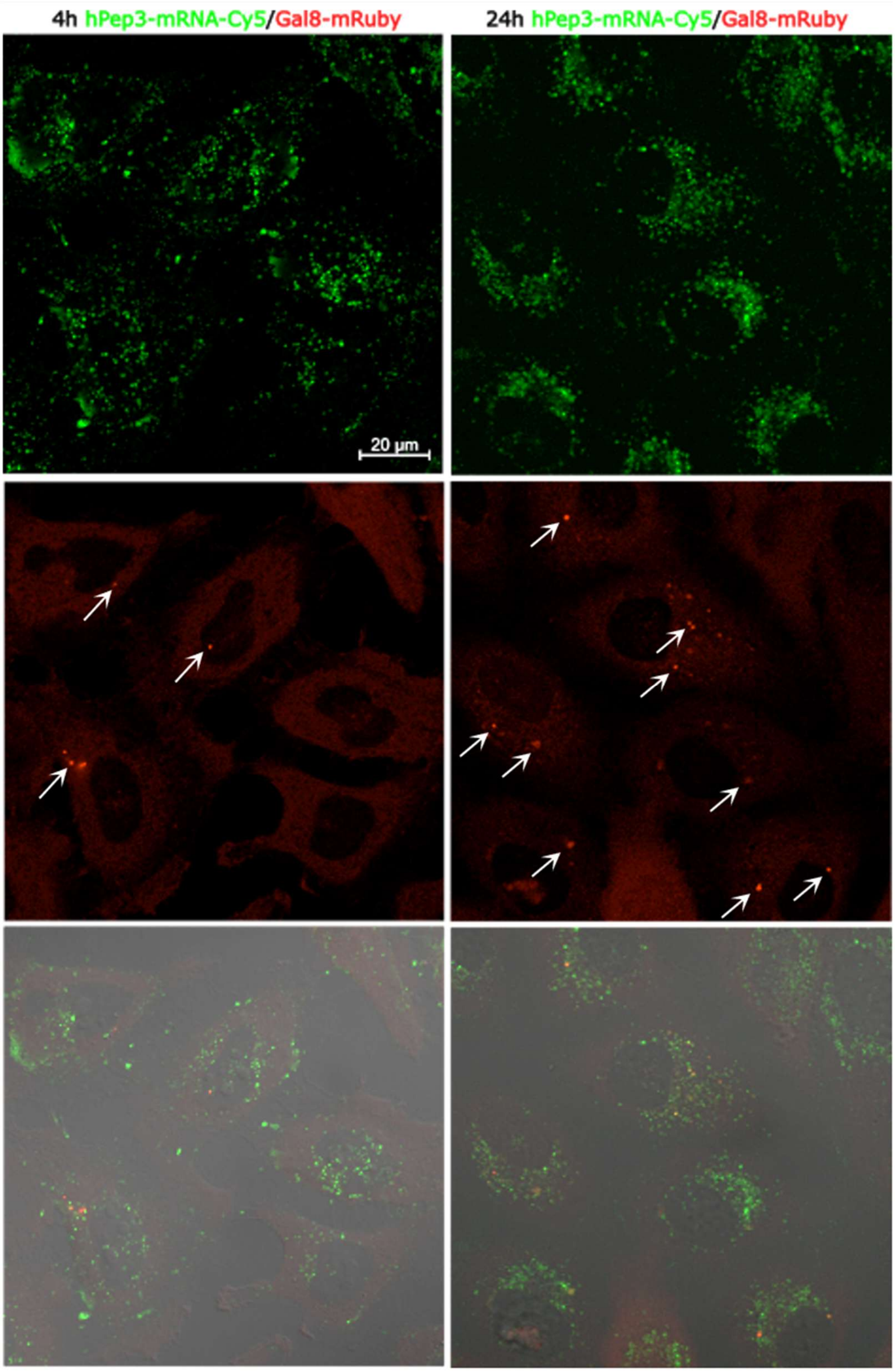


**Supplementary Figure 4. Endosomal release of hPep3/mRNA NPs.** Confocal microscopy of Gal8-mRuby expressing HeLa cells treated with Cy5-mRNA NPs at CR4 at the indicated time-points. Arrows indicate endosomal membrane disruption sites.

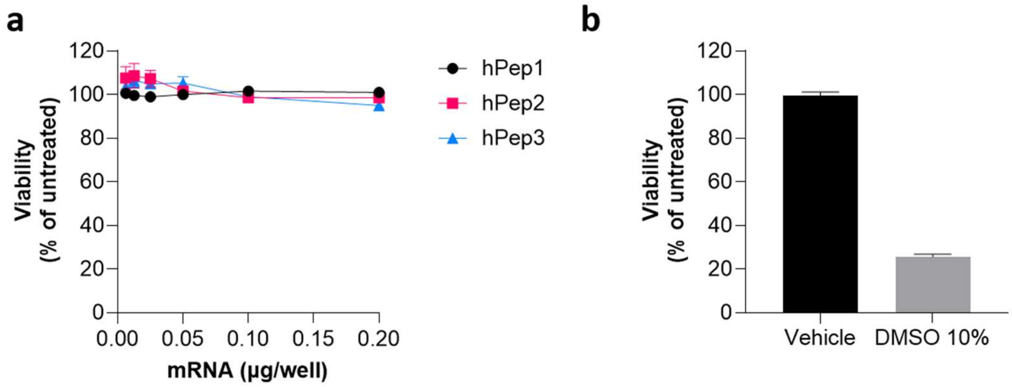


**Supplementary Figure 5. Dose-dependent evaluation of hPep/mRNA NPs on cell viability in A549 cells using WST-1 assay. a**, Viability of cells after 24-hour treatment with PNPs formulated at CR4, **b,** negative control formulation buffer (vehicle) and positive control (DMSO 10%). Data is presented as mean ± stdev (n=3).

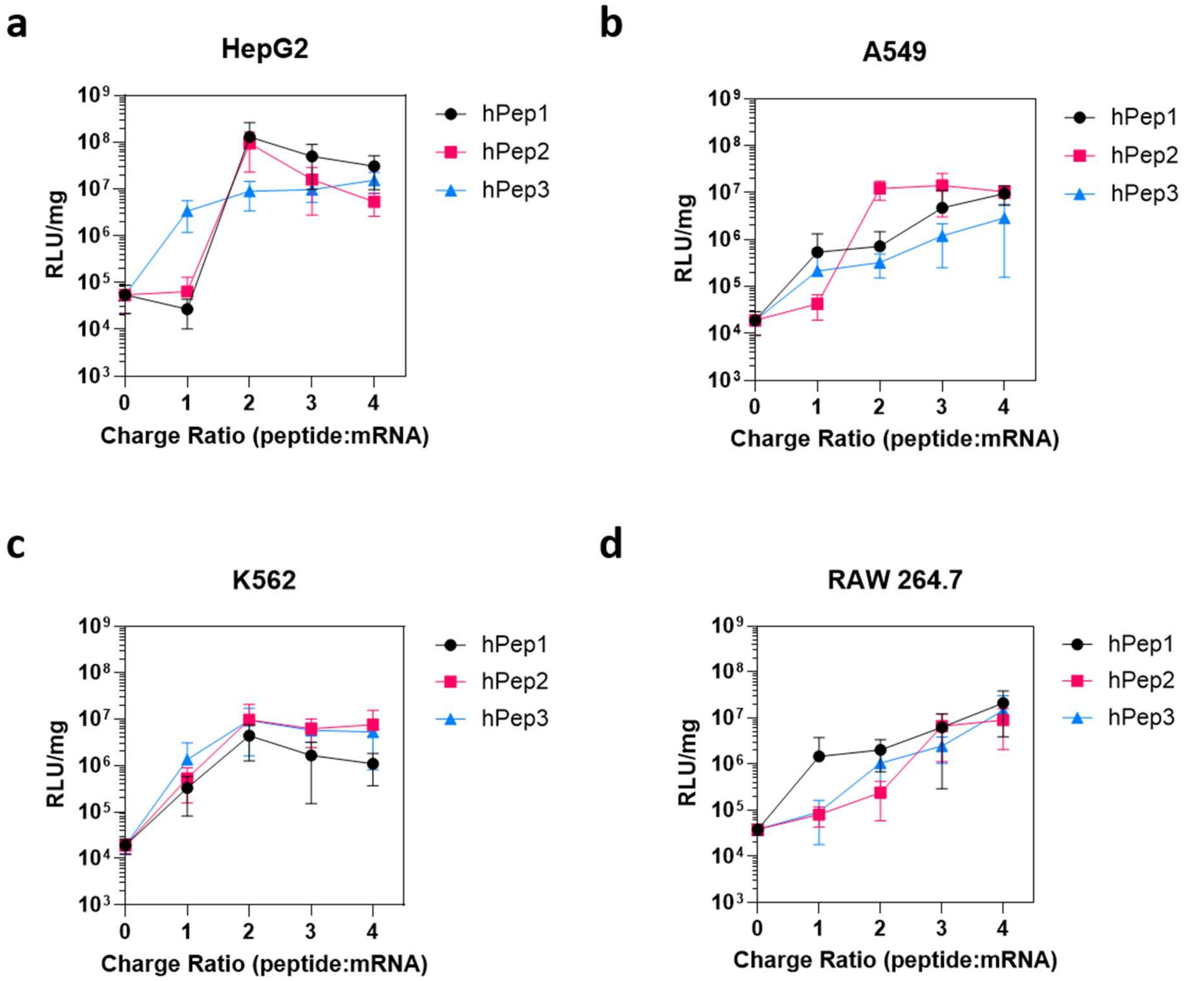


**Supplementary Figure 6. CR-dependent transfection efficacy of hPep/mRNA NPs in different cell lines. a-d**, Cell lines were treated for 24 h with hPep3/Fluc NPs formulated at different CRs and corresponding luciferase expressions are presented. Data is presented as mean ± stdev (n=3).

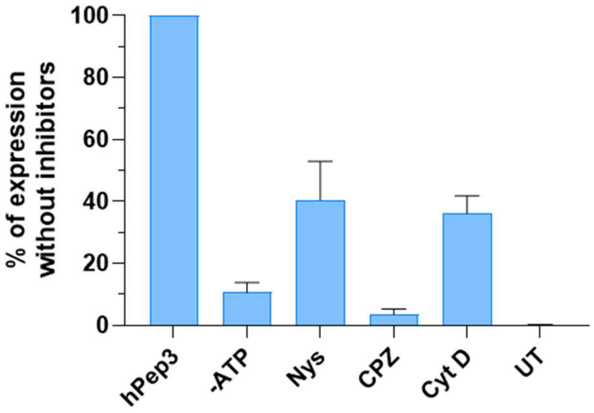


**Supplementary Figure 7. Effect of pharmacological endocytosis inhibition on the luciferase encoding mRNA delivery efficacy with hPep3/Fluc NPs in A549 cells.** Data is presented as mean ± SEM, n=3.

**Supplementary Table 1. Summary of all developed hPep analogs and their chemical properties**

| Name | Structure | Charge | Mw (g/mol) | α-helical content % | NP formation |
|---|---|---|---|---|---|
| **hPep1** | $NH_2$-LAKLAKA**{R8}**AKLLKA**{S5}**AKAL-$NH_2$ | +6 | 2040.4 | 14.9 | + |
| **hPep2** | $NH_2$-LAKLAKA**{R8}**AKLLKA**{S8}**AKAL-$NH_2$ | +6 | 2084.4 | 32.2 | + |
| **hPep3** | $NH_2$-L**{R8}**KLAKA**{R8}**AKLLKA**{S8}**AKAL-$NH_2$ | +6 | 2193.9 | 71.7 | + |
| hPep3-Ala, Ctrl | $NH_2$-L**A**KLAKA**A**AKLLKA**A**AKAL-$NH_2$ | +6 | 1863.5 | 0.0 | - |
| C8-hPep3 | Octanoyl-L**{R8}**KLAKA**{R8}**AKLLKA**{S8}**AKAL-$NH_2$ | +5 | 2320.4 | 15.3 | - |
| hPep3-OctG | $NH_2$-L**{OctG}**KLAKA**{OctG}**AKLLKA**{OctG}**AKAL-$NH_2$ | +6 | 2157.4 | 57.7 | + |
| hPep3-2Ado | $NH_2$-L**{2Ado}**KLAKA**{2Ado}**AKLLKA**{2Ado}**AKAL-$NH_2$ | +6 | 2241.4 | 64.0 | + |
| hPep3-K(C8:0) | $NH_2$-L**{K(C8:0)}**KLAKA**{K(C8:0)}**AKLLKA**{K(C8:0)}**AKAL-$NH_2$ | +6 | 2414.3 | 28.6 | + |
| hPep3-K(C14:0) | $NH_2$-L**{K(C14:0)}**KLAKA**{K(C14:0)}**AKLLKA**{K(C14:0)}**AKAL-$NH_2$ | +6 | 2666.8 | 72.8 | + |
| hPep3-K(C18:0) | $NH_2$-L**{K(C18:0)}**KLAKA**{K(C18:0)}**AKLLKA**{K(C18:0)}**AKAL-$NH_2$ | +6 | 2835.1 | 69.4 | + |
| hPep3-YYY | $NH_2$-L**Y**KLAKA**Y**AKLLKA**Y**AKAL-$NH_2$ | +6 | 2139.8 | 0.0 | - |
| C12-hPep3-YYY | Lauryl-L**Y**KLAKA**Y**AKLLKA**Y**AKAL-$NH_2$ | +5 | 2322.1 | 4.2 | + |
| hPep3-YFF | $NH_2$-L**Y**KLAKA**F**AKLLKA**F**AKAL-$NH_2$ | +6 | 2107.8 | 0.0 | - |
| C12-hPep3-YFF | Lauryl-L**Y**KLAKA**F**AKLLKA**F**AKAL-$NH_2$ | +5 | 2290.1 | 11.8 | + |
| hPep3-YWW | $NH_2$-L**Y**KLAKA**W**AKLLKA**W**AKAL-$NH_2$ | +6 | 2185.9 | 0.0 | - |
| C12-hPep3-YWW | Lauryl-L**Y**KLAKA**W**AKLLKA**W**AKAL-$NH_2$ | +5 | 2368.2 | 28.4 | + |

{R8}= (R)-2-(7-octenyl)alanine
{S5} = (S)-2-(4-pentenyl)alanine
{S8} = (S)-2-(7-octenyl)alanine
{OctG} = L-octylglycine
{2Ado} = (S)-2-amino-dodecanoic acid
{K(C8:0)} = N(6)-(octanoyl)lysine
{K(C14:0)} = N(6)-(myristoyl)lysine
{K(C18:0)} = N(6)-(stearoyl)lysine

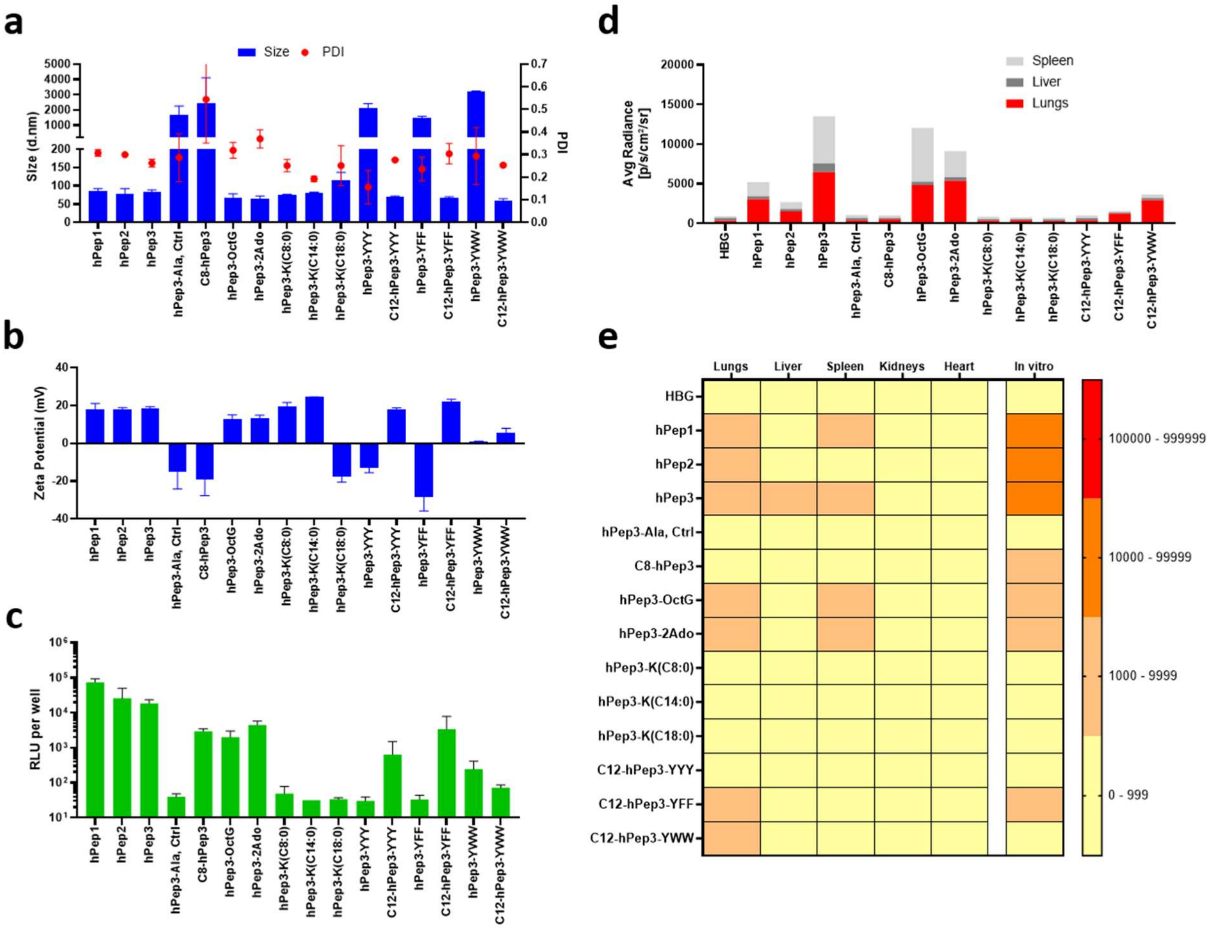


**Supplementary Figure 8. Characterization of peptide/mRNA NPs with hPep analogs and their efficacy in vitro and in vivo. a-b,** Size and zeta potential of hPep/mRNA formulations at CR4 as measured by DLS. **c**, A549 cells were treated for 24 h with hPep/Fluc-mRNA NPs formulated at CR4, and corresponding luciferase expressions are presented. **d,** For in vivo evaluation Balb/c mice (n=3 per group) were intravenously administered with hPep/mRNA NPs formulated at CR4 at a final dose of 1 mg/kg. After 24 h luciferase expression was quantified from dissected tissues with IVIS. **e,** Summary of in vivo and in vitro data. Data is presented as mean ± stdev (n=3).

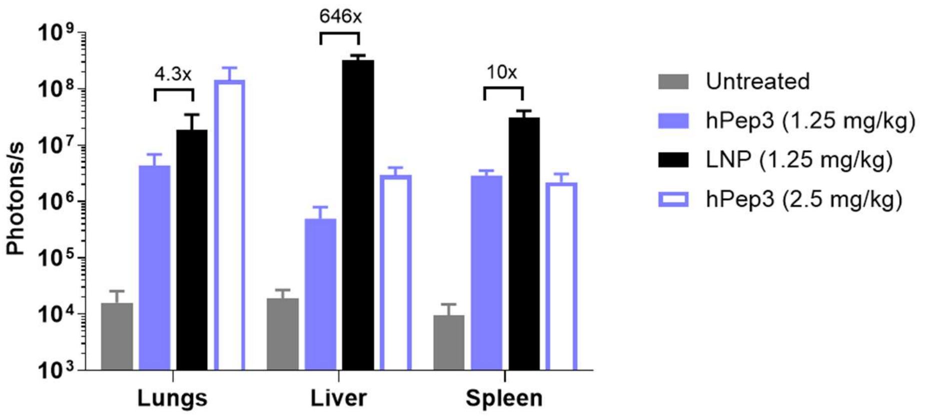


**Supplementary Figure 9. In vivo efficacy comparison between hPep3/mRNA NPs and MC3 LNPs.** Animals were treated with bulk mixed hPep3/mRNA NPs and microfluidically formulated LNPs at 1.25 mg/kg mRNA dose. After 24 h tissues were dissected and measured under IVIS. Data is presented as mean ± stdev (n=3). Values represent the signal fold increase of LNPs over hPep3 NPs.

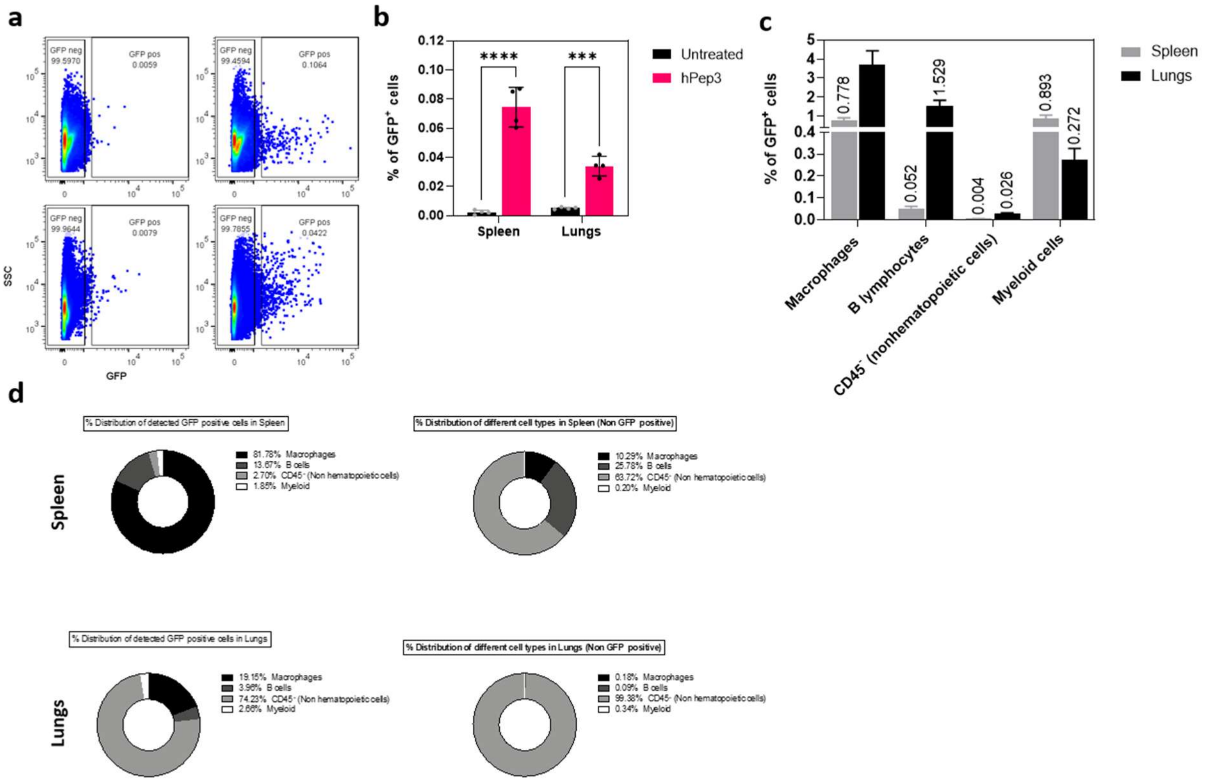


**Supplementary Figure 10. Target cell type identification of hPep3/EGFP-mRNA PNPs in vivo. a,** Mice (n=3 animals per group) were treated with hPep3/EGFP mRNA NPs i.v., followed by flow cytometry analysis of the tissues 24 hours post-injection. **b,** Quantification of the overall EGFP expression in different tissues by FACS. **c, d,** Cell type-specific expression of EGFP mRNA, verified 24 hours after injection of hPep3/mRNA NPs using FACS. (n=3 animals per group). Data is presented as mean ± stdev. Statistics: two-way ANOVA.

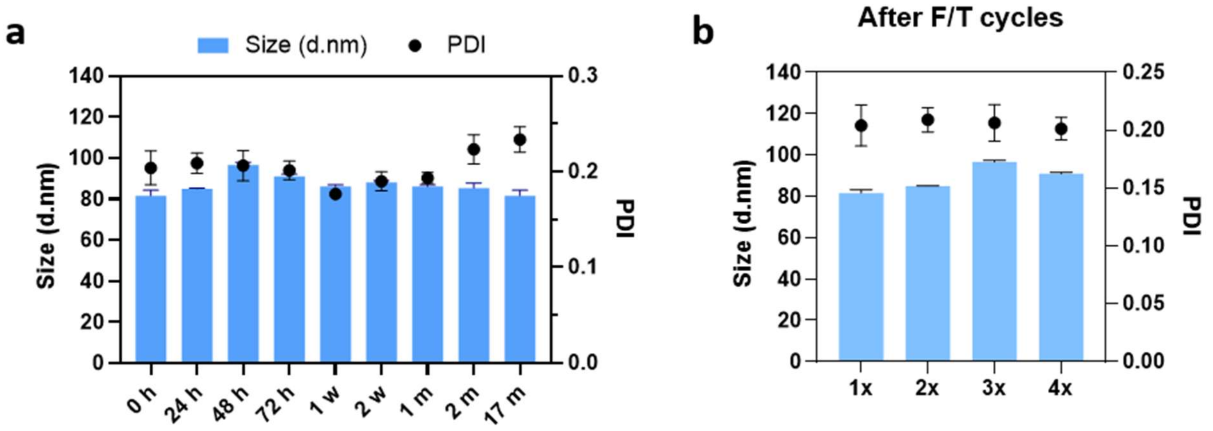


**Supplementary Figure 11. Short- and long-term stability of microfluidically formed hPep/mRNA formulations in cryostorage. a,** Formulation was carried out with NanoAssemblr™ Blaze™ instrument in combination with NxGen 500 cartridge with hEPO mRNA and stored at -80°C over time and after subjecting to **b,** freeze-thaw cycles. DLS data is presented as mean ± stdev.

**Supplementary Table 2. Formulation process parameters for NHP trial formulations.**

| Parameter | Description |
|---|---|
| Instrument | NanoAssemblr™ Blaze™, Cytiva |
| Mixer type | NxGen 500 |
| hPep3/mRNA w/w | 8.0 |
| hPep3 in water | 2.4 mg/ml |
| mRNA in water | 0.3 mg/ml |
| Flow rate ratio (FRR) | 1:1 |
| Total flow rate (TFR) | 100 ml/min |
| Output mRNA concentration | 0.15 mg/ml |
| Tangential flow filtration (TFF) settings | **FLuc NPs**<br>150 ml/min, transmembrane pressure (TMP) 4 psi, shear rate ~5500 $sec^{-1}$ through a D02-E030-05-N (115 $cm^2$) column with L/S 16 tubing.<br>**hEPO NPs**<br>150 ml/min, TMP 4 psi, shear rate ~5500 $sec^{-1}$ through a S02-E030-05-N (790 $cm^2$) column with L/S 17 tubing. |
| mRNA target concentration after TFF | ≥1.4 mg/ml |
| Dilution | 1:1 with 2X HEPES buffered sucrose (SH) |
| Sterile filtration | 0.22 µm PES, Minisart®, 25 mm filter |
| mRNA and encapsulation efficiency quantification | RiboGreen assay with heparin treatment |
| mRNA dilution to target concentration | 0.6 mg/ml, diluted with sterile 1X SH buffer (10 mM HEPES, 10% w/v sucrose, pH7.4) |
| Aliquotation and storage | 5 ml aliquots stored at -80 °C |
| NP characterization | DLS, Cryo-EM, endotoxin testing, peptide concentration with BCA assay |

**Supplementary Table 3. Additional NHP trial formulation parameters**

| | hEPO | FLuc |
|---|---|---|
| Peptide/mRNA w/w (PNI nanodrop_diluted stocks) ±SD* | 9,73±0,4 | 6,50±1,8 |
| Peptide/mRNA w/w (our calc) ** | 5,85 | 3,85 |
| Starting mRNA amount | 190 mg | 40 mg |
| mRNA amount in formulation | 148 mg | 35 mg |
| Yield (mRNA) | 78% | 87% |

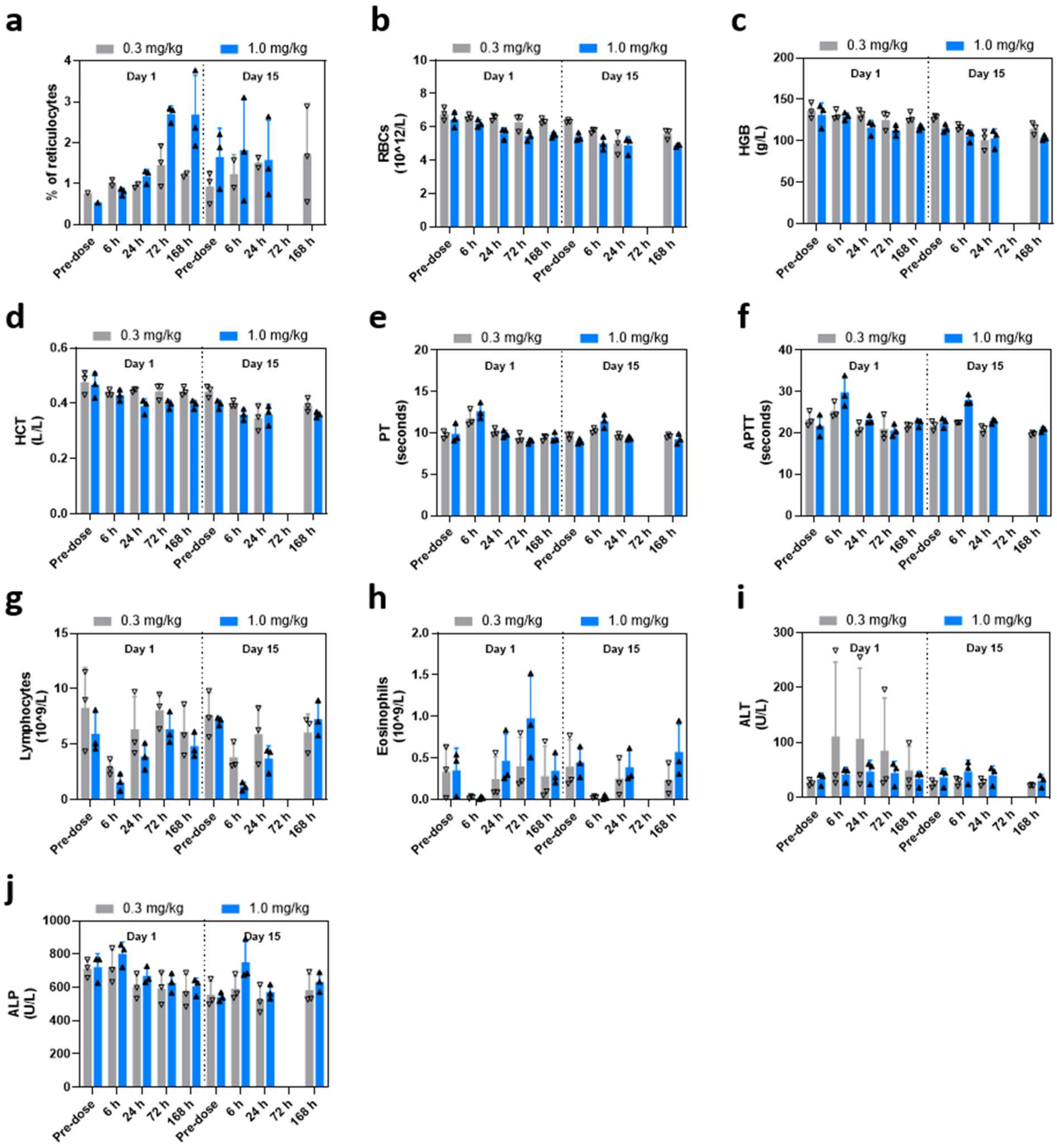


**Supplementary Figure 12. Clinical parameters of repeated administration hPep3/hEPO mRNA NPs in NHPs. a,** Blood reticulocytes percentage, **b,** red blood cell count, **c,** hemoglobin concentration, **d,** hematocrit level, **e,** prothrombin time, **f,** activated partial thromboplastin time, **g,** lymphocyte count, **h,** eosinophil count, **i,** alanine transaminase concentration and **j,** alkaline phosphatase concentration. Data is presented as mean ± stdev.

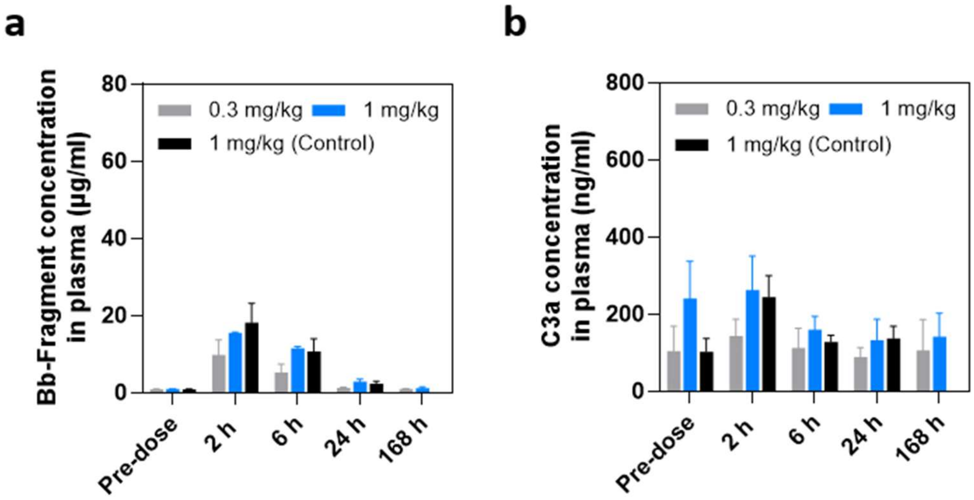


**Supplementary Figure 13. Complement activation after single administration of hPep3/hEPO mRNA and hPep3/Control mRNA formulations in NHPs is mild and transient. a,** Bb-fragment and **b,** C3a factor concentrations were measured from plasma at noted time-points, Data is presented as mean ± stdev (n=3 animals per group).
